# A microbiota-derived protease links phage susceptibility to host epithelial responses

**DOI:** 10.64898/2026.09.26.754296

**Authors:** Ning Qu, Marlène S Birk, Thomas C A Hitch, Charlotte De Rudder, Beatriz Monteiro, Musthofa Kemal Bandu, Marla Gaissmaier, Lina Michel, Monica Steffi Matchado, Carlos Geert Pieter Voogdt, Aicha Kriaa, Nicole S Treichel, Bjoern O. Schroeder, Eva Miriam Buhl, Christian Preisinger, Athanasios Typas, Bärbel Stecher, Paul Wilmes, Thomas Clavel, Joel Selkrig

## Abstract

Bacteriophages are major ecological drivers of gut microbial ecology, yet whether bacterial mechanisms that determine phage susceptibility have consequences for the mammalian host remains poorly understood. Here, we identify dipeptidyl peptidase 11 (Dpp11a), the predominant active serine protease of the prevalent gut commensal *Phocaeicola vulgatus*, as an unexpected bacterial defence factor. Dpp11a protects against environmental proteases and confers resistance to bacteriophage infection. Metatranscriptomic analyses further reveal increased expression of both *dpp11a* and *P. vulgatus*-associated phage transcripts in ulcerative colitis stool samples, indicating that both components of this interaction are transcriptionally active in disease-associated human microbiomes. Using the microfluidic gut-on-a-chip co-culture model HuMiX, we show that the absence of Dpp11 is accompanied by altered epithelial tight-junction remodelling during phage-bacterial infection. Together, our findings reveal that the consequences of bacterial phage defence can extend beyond phage-bacterium interactions to the mammalian epithelium.

## Introduction

The intestinal lumen is one of the most proteolytically active microbial ecosystems in the body, shaped by host digestive enzymes together with proteases produced by the gut microbiota and intestinal epithelium^1–10^. Functional metagenomic studies have established that microbial proteases are abundant throughout the human gut and represent a major contributor to intestinal proteolytic activity^4–7^. Beyond their established roles in peptide metabolism, microbiome-derived proteases are increasingly recognized as important modulators of host physiology and disease^4,6,11–14^. However, despite their abundance and growing biomedical relevance, the ecological functions of these enzymes for the bacteria that produce them remain poorly understood.

Inflammatory bowel disease (IBD) provides a unique ecological setting in which gut bacteria encounter multiple concurrent selective pressures. Intestinal inflammation is associated with elevated luminal trypsin-like activity, accumulation of neutrophil-derived serine proteases, and extensive remodeling of intestinal protease homeostasis^1,7,9,15^. At the same time, ulcerative colitis (UC), a major subtype of IBD, is characterized by marked alterations of the gut virome, including expansion of tailed bacteriophages, disruption of bacterial community structure, and the presence of temperate phages capable of infecting *Phocaeicola vulgatus* (previously *Bacteroides vulgatus*) and related *Bacteroides* species^16–22^. Together, these observations suggest that gut bacteria must simultaneously withstand elevated extracellular proteolytic stress and bacteriophage predation during intestinal inflammation. Yet whether shared bacterial mechanisms confer resistance to both selective pressures remains unknown.

*P. vulgatus* is not only one of the most abundant members of the healthy human gut microbiota, it also plays an important role in UC pathobiology. Strain-level serine protease activity of *P. vulgatus* correlates with endoscopic and histological disease severity in UC patients, and the transfer of protease-rich patient faecal communities into germ-free mice induces protease-dependent colitis^13^. *P. vulgatus* strains isolated from UC patients show enhanced adherence to intestinal epithelium relative to strains from healthy individuals ^23^, and defined *P. vulgatus* strains are sufficient to induce colitis in genetically susceptible hosts, including monoassociated HLA-B27 transgenic rats ^24,25^ and, in a host-genotype-dependent manner in murine IBD models ^26,27^. In parallel, *P. vulgatus* and closely related *Bacteroides* species are among the principal bacterial hosts of the tailed bacteriophages that expand during UC^16,19–22,28^. *P. vulgatus* therefore occupies a position in which elevated luminal proteolysis and intensified phage predation converge on a single, clinically implicated taxon, thereby making the mechanisms by which it withstands these pressures directly relevant to disease.

Among bacterial serine peptidases, dipeptidyl peptidases (DPPs) are a highly conserved family of serine proteases best known for liberating dipeptides from oligopeptides during nutrient acquisition ^29^. More recently, microbial DPPs have emerged as important mediators of host-microbiome interactions, with bacterial Dpp4-like enzymes shown to function as host-microbe isozymes that process incretin hormones and other bioactive peptides, thereby influencing host metabolism and therapeutic responses^30–32^. Despite this growing appreciation of their host-directed activities, it remains unknown whether DPPs contribute more broadly to bacterial fitness and ecological adaptation within the intestinal ecosystem.

Here, we identify Dpp11a as the dominant active serine protease produced by one of the most common and abundant gut microbiome species, *P. vulgatus,* and demonstrate that it has a dual defense role in protecting against both extracellular proteolytic stress and bacteriophage infection. Integrating activity-based proteomics, bacterial genetics, microfluidic gut-on-a-chip co-culture in HuMiX, and human metatranscriptomic analyses, we further show that this defense pathway is likely activated in UC and modulates host epithelial responses during phage infection. Together, our findings reveal an unexpected ecological function for a highly abundant gut microbial peptidase and implicate microbial phage defense as a previously unrecognized link between microbiome ecology and host physiology.

## Results

### Serine proteases dominate the human gut microbiome protease repertoire

To investigate the diversity of proteolytic repertoires across human gut bacterial taxa, we analysed MEROPS^33^ (the Peptidase Database) annotations for 31 fully sequenced representative bacterial species isolated from healthy human gut microbiomes ^34^ (see **Table S1** for complete species list). Of all predicted protease-encoding genes across these species, serine proteases represented the most common protease family (39.33%), followed by metalloproteases (29.47%), cysteine (17.26%), aspartic (7.7 %), glutamic (2.16%) and asparagine (1.8%) proteases (**Fig. 1a**). When species are ranked proportionally to their relative abundance in healthy human feces^35–37^, serine proteases account for the largest proportion of the predicted microbiome protease repertoire overall (**Fig. 1a, Fig. S1a**). Accordingly, *P. vulgatus* was one of the largest contributors to the serine protease family among individual taxa.

**Figure 1.**
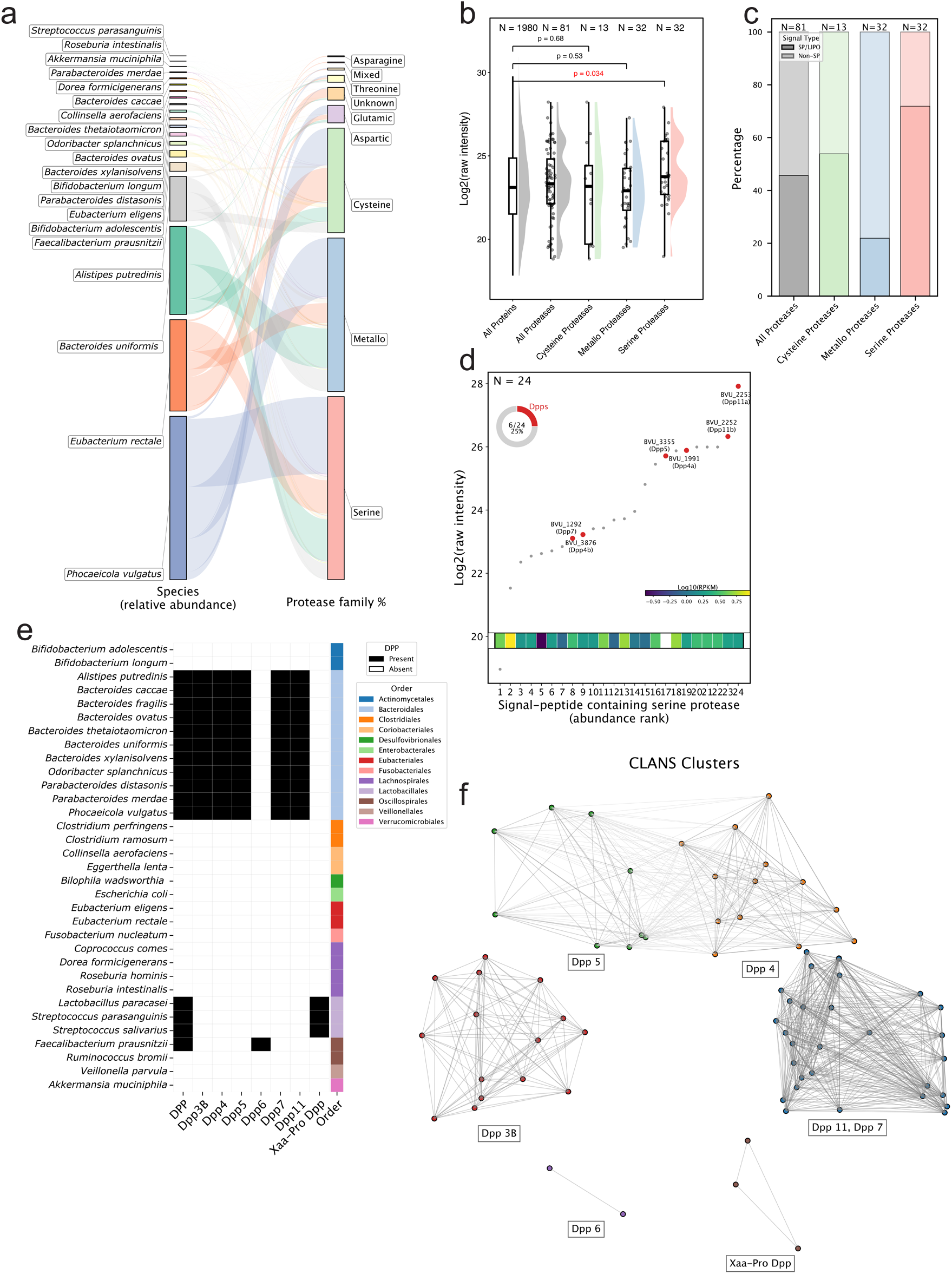
Dipeptidyl peptidases are an abundant and prevalent class of serine proteases in the human gut microbiome. **a**) Abundance-weighted distribution of bacterial protease families in the human gut microbiota (see Table S1 for strain list). Bacterial species are shown on the left and protease families on the right. Species bar heights reflect the square of their median relative abundance across samples. Ribbons indicate the protease families encoded by each species, with ribbon widths representing the abundance-weighted contribution of each species to a given protease family (species abundance × fraction of encoded proteases belonging to that family). Protease family bar heights show the cumulative abundance-weighted representation of each family across all analysed species. Ribbons

Given the large repertoire of serine protease-encoding genes in *P. vulgatus*, its abundance in the healthy human gut microbiome ^34,38,39^, and the association of *P. vulgatus*-derived serine proteases with UC severity^13^, we next used LC-MS/MS to identify the most abundant proteases when grown *in vitro*. While the number of detected serine- and metallo-proteases was identical (n = 32), serine proteases were significantly more abundant compared to the rest of the proteome (p = 0.034 two-sided Mann-Whitney U test), whereas metallo- and cysteine-proteases showed no significant difference (**Fig. 1b**). These data demonstrate that serine proteases are not only the most frequently encoded protease family, but also a highly abundant protease class produced by *P. vulgatus*, suggesting an important role in gut microbial ecology.

As secreted gut microbial proteases can shape microbial communities and their behaviour ^40,41^, as well as direct interactions with the host^14,42^, we next examined for the presence of N-terminal signal peptides in proteases expressed by *P. vulgatus*. We observed that ∼75% of *P. vulgatus* serine proteases (24/32 serine proteases) are predicted to encode signal peptides by SignalP^43^, indicating serine-proteases are typically targeted to the cell envelope and/or extracellular milieu in this species (**Fig. 1c**). In contrast, only 20% of metalloproteases and 53% of cysteine proteases, respectively, encode putative signal peptides. These observations are in line with previous reports whereby serine proteases were frequently detected in spent culture supernatants of *P. vulgatus* and related species e.g. *Phocaeicola dorei* and *Bacteroides thetaiotaomicron*^13^, indicating *P. vulgatus* serine proteases play key roles in the cell envelope and/or the extracellular space. are coloured by bacterial species and protease family bars by catalytic class. Protease annotations were obtained from the MEROPS database^89^. **b**) Raw LC-MS/MS intensity values of whole cell proteomes of *Phocaeicola vulgatus*^ATCC 8482^ grown in monoculture, where proteins are separated into indicated protease Family categories. Boxes represent the interquartile range (IQR), with the center line indicating the median. Whiskers indicate the minimum and maximum values. Individual observations are shown as grey dots, and violin plots depict the data distribution, *n* denotes the number of proteins in each group. Combined data form three independent experiments. A two-sided Mann-Whitney U test was used to test significance. **c**) Fraction of each protease Family predicted to harbor an n-terminal signal peptide (SP) or Lipo-signal peptide (LIPO) in *P. vulgatus*^ATCC 8482^. *n* refers to the number of proteins in each group (total). **d**) scatterplot of putative serine-protease-encoding proteins encoding a signal peptide from

*P. vulgatus*^ATCC 8482^ ranked according to cellular abundance as a function of raw intensities, the most abundant of which is labelled in red: dipeptidyl-peptidase 11; Dpp11 (BVU_2253). Inset heatmap indicates Log_10_ reads per kilobase million (RPKM_Log10_) transcripts for each gene obtained by searching candidate genes from publicly available metatranscriptomics datasets^42^. The fraction of *P. vulgatus*^ATCC 8482^ signal-peptide-encoding serine proteases belonging to the DPP class are shown in the upper left. **e**) phylogenetic analysis of dipeptidyl-peptidase (DPP) encoding genes across diverse commensal gut microbiome species. Presence and absence of DPP homologs across 31 representative gut-associated bacterial species from (a) were determined using MEROPS-based annotations. **f**) CLuster ANalysis of Sequences (CLANS^86^) depicts an all-against-all pairwise BLAST analysis of DPP protein sequences. Higher data point proximity reflects greater sequence similarity. Nodes correspond to DPPs in a given species. Node edges are drawn between similar sequences based on a *P* value cutoff ≤1e^−6^.

Of the 24 signal-peptide encoding serine proteases expressed in *P. vulgatus*, 25% (6/24 proteins) belonged to the pharmaceutically relevant enzyme class referred to as dipeptidyl peptidases (DPPs) (**Fig. 1d**). Two DPPs ranked among the most highly abundant serine proteases in the *P. vulgatus* proteome (**Fig. 1d**), namely BVU_2253 and BVU_2252, herein referred to as Dpp11a and Dpp11b (according to MEROPS peptidase database nomenclature), respectively. Moreover, evidence for the expression of these genes at the transcript level from human fecal microbiome samples obtained from Lloyd-Price *et al* ^42^ indicate the genes encoding these corresponding proteases are indeed also expressed in the human gut (**Fig. 1d**, inset). These results demonstrate serine proteases likely play an important role in the *P. vulgatus* cell envelope and/or extracellular space, of which DPPs constitute a significant fraction of expressed serine proteases and are among the most abundant.

Although DPPs have been extensively characterized in oral pathogens for their role in peptide scavenging and nutrient acquisition^44^, they have only recently emerged as important mediators of microbiome-host interactions^30–32^. We therefore asked how broadly DPP-encoding genes are distributed across representative members of the human gut microbiota. We found DPPs were strongly conserved among members of the order *Bacteroidales*, with more limited representation in other taxa (**Fig. 1e**). Dpp3B, Dpp4, Dpp5, Dpp7 and Dpp11 were restricted to *Bacteroides*, whereas Dpp6 and Xaa-Pro-type DPPs were more commonly detected outside this genus. Sequence similarity analysis by CLANS further resolved these enzymes into distinct clusters: Dpp3B, Dpp6 and Xaa-Pro-type DPPs formed separate groups, whereas Dpp4 and Dpp5, and Dpp7 and Dpp11, clustered together, respectively (**Fig. 1f**). Their grouping was additionally supported by structural similarity (**Fig. S1b-c**), consistent with evolutionary divergence accompanied by subgroup-specific functional specialization. Together, these data identify DPPs as a diverse and prevalent subgroup of serine proteases in *Bacteroidales*.

### Dpp11a is the most abundant and active serine hydrolase in *P. vulgatus*

To identify proteolytically active serine proteases in *P. vulgatus*, we treated cell pellets with the activity-based probe TAMRA-fluorophosphonate (TAMRA-FP), which covalently labels the catalytic serine of active serine hydrolases. Whole cell lysates revealed several *P. vulgatus* proteins formed TAMRA-FP adducts of diverse molecular weights and intensities, with particularly prominent reactivity with proteins between 80-90 kDa (**Fig. 2a**). TAMRA-FP specificity was verified by the absence of these bands in solvent control samples. To further identify FP-reacting serine hydrolases, we performed affinity-based LC-MS/MS using Biotin-FP as bait on bacterial cells. We identified 28 significantly enriched proteins (> 2.0 Log2, ≤ 0.05 FDR, Benjamini-Hochberg adjusted) compared to background controls (**Fig. 2b**). As expected, the predicted enzyme functions of these significantly enriched proteins related to serine-type peptidases, dipeptidyl peptidase and serine-type amino peptidase activity (≤ 0.05 FDR, Benjamini-Hochberg adjusted) (**Fig. 2c**), confirming the specificity of the pulldown reaction for serine hydrolases, including serine peptidases. Moreover, we further validated the specificity of the affinity-purification reaction by determining which proteins could be out-competed by first occluding reactive-serine residues by TAMRA-FP (pre-blocking) immediately prior to Biotin-FP affinity purification LC-MS/MS. This revealed 15/31 enriched proteins (∼48%) were outcompeted by TAMRA-FP (competitive index ≤ -2 Log2), thereby providing strong evidence these proteins selectively interact with FP (**Fig. 2d**).

**Figure 2.**
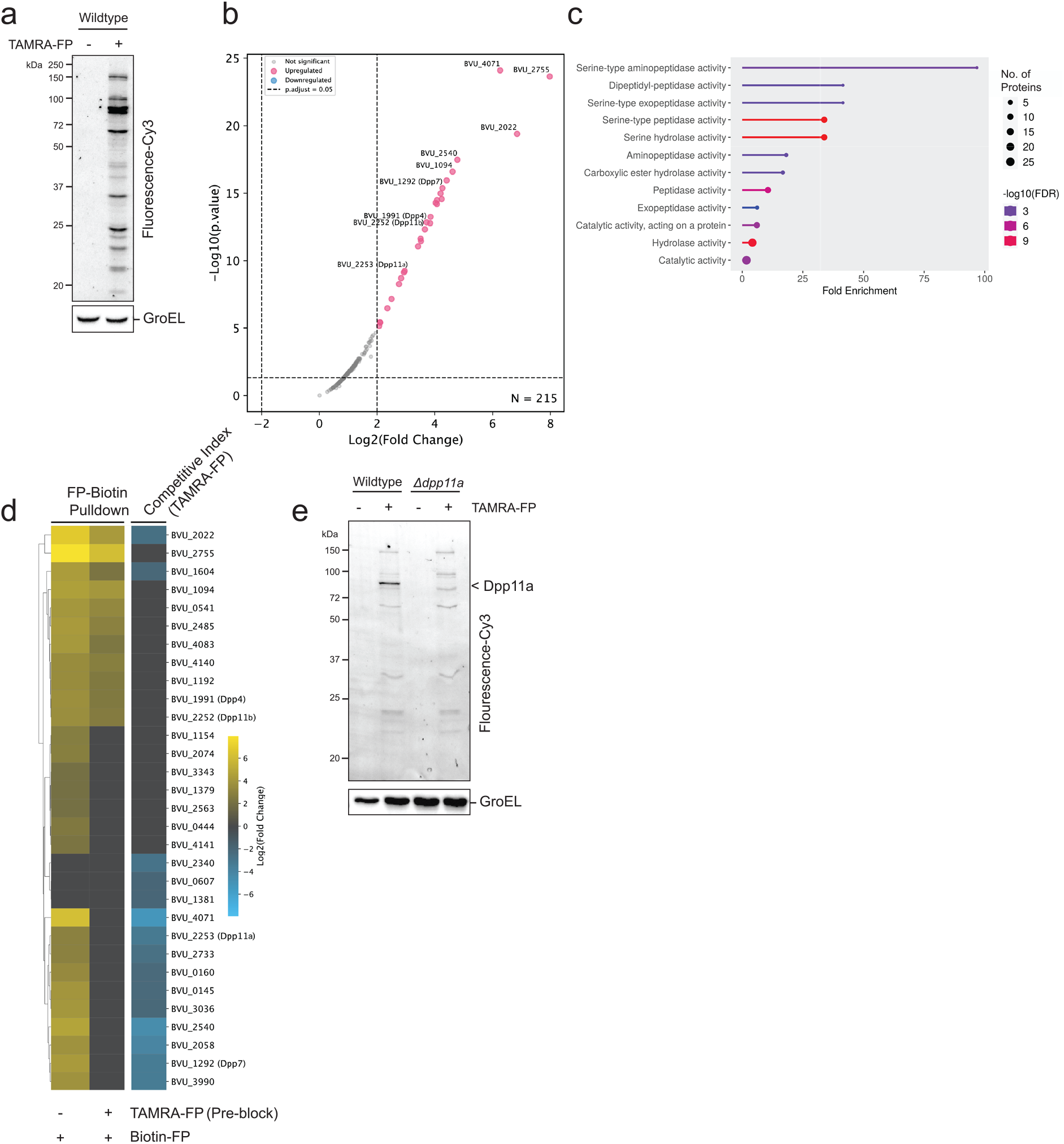
Dpp11a is a dominant serine protease of *P. vulgatus*. **a**) Intact exponentially growing *P. vulgatus* typed strain ATCC 8482 (*P. vulgatus* ^ATCC 8482^) grown in nutrient-rich mGAM media, reacted with 1 µM TAMRA-FP Serine Hydrolase activity Probe, proteins were separated by SDS-PAGE and in-gel visualisation of TAMRA-FP reactive proteins was performed by Cy-3 fluorescence detection (Ex/Em: 550 nm/570 nm). The same gel was then subjected to immunoblot analysis using α-GroEL (cytoplasmic) as a sample loading control. Representative image of three independent experiments (replicates in Supplementary Data). **b**) Intact *P. vulgatus*^ATCC 8482^ was reacted with 1 µM Biotin-FP, reactive proteins were enriched using streptavidin beads and eluted by on-bead trypsin digestion. Eluted peptides were labelled with TMT isobaric tags, measured by LC-MS/MS and compared to solvent control samples. Significantly enriched proteins passing Log_2_ Fold Change (FC) ≥ 2.0 and -log_10_ *P*-value of ≤ 0.05 (Benjamini-Hochberg adjusted) are plotted as red datapoints. Combined data from three independent experiments. **c**) GO enrichment analysis was performed by ShinyGO 0.85.1 Tool with default parameters using proteins from (b) passing indicated significance thresholds. **d**) Heatmap of proteins enriched using Biotin-FP coupled to LC/MS/MS and passing significance thresholds as described in (b) compared to the same samples pre-blocked with 1 µM TAMRA-FP. Relative differences of enrichment of the indicated proteins was calculated by subtracting Biotin-FP Log_2_ enrichment from TAMRA-FP Log_2_ pre-blocked samples and expressed as the Competitive Index. Combined data from three independent experiments. Corresponding Data is located in supplementary data. **e**). Intact *P. vulgatus*^ATCC 8482^ cells and an isogenic *Δdpp11* (BVU_2253) deletion mutant was reacted with TAMRA-FP and analysed as described in (a). Representative data of three independent experiments (replicates in Supplementary Data).

To identify the serine protease responsible for the dominant TAMRA-FP-reactive adducts, we tested a panel of *P. vulgatus* transposon mutants carrying insertions in candidate protease-encoding genes for the loss of TAMRA-FP reactivity. Strikingly, transposon disruption of Dpp11a abolished the predominant ∼80 kDa TAMRA-FP adduct (**Fig. 2e**, **Fig. S2a**), directly implicating Dpp11a as the major active serine protease. Consistent with this assignment, Dpp11a labeling was highly sensitive to the irreversible serine protease inhibitor phenylmethylsulfonyl fluoride (PMSF) (**Fig. S2b**). Together, these findings establish Dpp11a as the predominant catalytically active serine protease expressed by *P. vulgatus*.

### Dpp11a localises to the periplasm and extracellular space

Having identified Dpp11a as the major active serine protease in *P. vulgatus*, we next asked whether its subcellular localization was consistent with a role in protecting the bacterium from extracellular challenges. Although most *Bacteroidales* DPPs are thought to reside within the periplasm^44^, some DPPs have been reported to be secreted extracellularly ^13,29^. To determine whether Dpp11a is positioned to interact with extracellular substrates, we examined its subcellular distribution by coupling cell fractionation with TAMRA-FP labeling in wild-type and an isogenic Δ*dpp11a* strain. The majority of Dpp11a activity was associated with intact cells (**Fig. 3a**), although approximately 20% of the activity was consistently detected in unconcentrated cell-free supernatants from exponentially growing cultures (**Fig. 3a,b**). Importantly, this extracellular activity was not attributable to cell lysis, as the cytoplasmic marker GroEL, used as a loading control, remained undetectable in cell-free supernatants even after prolonged exposure. We then asked whether cell-bound Dpp11a localised to the periplasm by subjecting *P. vulgatus* overexpressing C-terminally HA-tagged Dpp11a (Dpp11a-HA) to osmotic shock which selectively releases soluble periplasmic proteins^45^. Dpp11a-HA was released into the osmotic shock extract, whereas the soluble cytoplasmic marker GroEL was not, confirming that the inner membrane remained intact and that selective extraction of the cell envelope had occurred (**Fig. S2c**). To determine whether the released protein was membrane-associated or soluble periplasmic, osmotic shock extracts were further fractionated by ultracentrifugation. Dpp11a was recovered exclusively in the soluble fraction, together indicating a soluble periplasmic localisation. Finally, immunogold transmission electron microscopy independently confirmed cell-envelope localization of cell-associated Dpp11a-FLAG, revealing specific gold labelling surrounding bacterial cells, which was absent in the wild-type controls (**Fig. 3c**). Together, these complementary approaches establish that Dpp11a is predominantly a soluble periplasmic protease while also being released into the extracellular environment.

**Figure 3.**
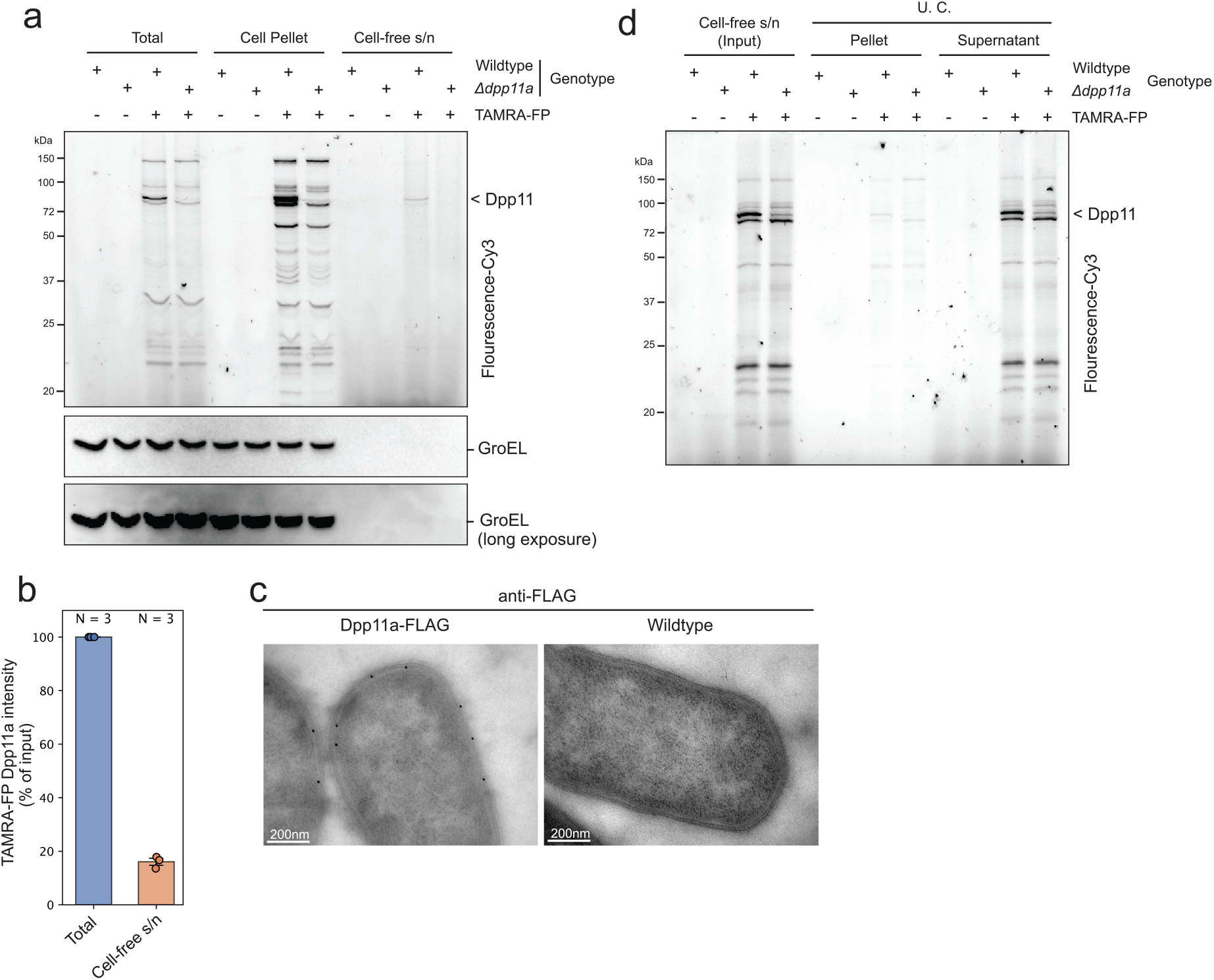
Dpp11a localizes to both the periplasm and the extracellular milieu while retaining enzymatic activity. **a**) Exponentially growing *P. vulgatus*^ATCC 8482^ and an isogenic Δ*dpp11a* mutant were separated into total culture, cell pellet, and cell-free supernatant (s/n) fractions by centrifugation. Cell pellets were resuspended in PBS, and all fractions were labelled with the activity-based probe TAMRA-FP and analysed by SDS-PAGE fluorescence scanning as described in Fig. 2a. The lower panels show α-GroEL immunoblots of the corresponding samples; the bottom blot is a longer exposure to detect GroEL in the cell-free supernatant. Representative of three independent experiments (replicates shown in Supplementary Data). **b**) Quantification of extracellular Dpp11a activity detected in cell-free supernatants, expressed as a percentage of the total input TAMRA-FP-labelled Dpp11a signal. Points represent independent biological replicates (N = 3); bars show mean ± s.e.m. **c**) Immunogold transmission electron microscopy using an α-FLAG antibody demonstrating cell envelope-associated Dpp11a-FLAG staining. Wild-type cells lacking the FLAG epitope served as a negative control. Scale bars, 200 nm. **d**) Cell-free supernatants from exponentially growing wild-type and Δ*dpp11a* cultures were concentrated (∼30-fold) using a 3-kDa molecular weight cut-off filter and separated into membrane-associated (pellet) and soluble (supernatant) fractions by ultracentrifugation. Fractions were labelled with TAMRA-FP and analysed as in Fig. 2a. Dpp11a activity was detected predominantly in the soluble extracellular fraction. Representative of three independent experiments (replicates shown in Supplementary Data).

The detection of extracellular Dpp11a prompted us to determine whether it is released as a freely soluble enzyme or packaged within outer membrane vesicles (OMVs), a common protein export mechanism in Gram-negative bacteria. Spent culture supernatants were further sub-fractionated by ultracentrifugation to separate soluble proteins from membrane-associated material followed by TAMRA-FP labeling (**Fig. 3d**). Dpp11a activity was recovered almost exclusively in the soluble fraction, demonstrating that extracellular Dpp11a is released predominantly as an active soluble protease rather than in association with OMVs.

### Dpp11a promotes defense against environmental proteases and phage attack

Given its dual localization to both the periplasm and extracellular environment, we hypothesized that Dpp11a may contribute to protection against extracellular enzymatic attacks^46^ from neighboring bacteria or the host. Although Dpp11a and the downstream protein Dpp11b share strong structural similarity with the *P. gingivalis* dipeptidyl peptidase Dpp11 (**Fig. S3a–c**), we wondered whether these proteins have diverged to perform distinct biological functions. In *P. gingivalis,* Dpp11 mutants were previously shown to display growth defects compared to isogenic wildtype strains^29^. Our growth curve analysis of *P. vulgatus* isolates containing transposon insertions in Dpp11a and Dpp11b revealed both isolates grow similarly to the wildtype strain in nutrient rich mGAM media (**Fig. S3d**). In contrast to Dpp11a, Dpp11b and Dpp7 (BVU_1292) exhibited a mild growth delay in semi-synthetic Varel-Bryant medium, suggesting that Dpp11b retains a role in nutrient acquisition under low-nutrient conditions, akin to *P. gingivalis* Dpp11 (**Fig. S3d**). Consistent with potential functional specialization following gene duplication, CLANS sequence similarity analysis resolved *P. vulgatus* Dpp11a and Dpp11b into distinct sub-clusters together with their respective orthologues from other *Bacteroidales*, supporting evolutionary divergence between the two paralogues (**Fig. S3e**). Together, these findings suggest that, despite their structural similarity, Dpp11a and Dpp11b may have evolved distinct cellular functions.

We next asked whether extracellular localization of Dpp11a confers a functional advantage by protecting *P. vulgatus* against proteolytic stress. This question is particularly relevant in inflammatory bowel disease (IBD), where elevated luminal serine protease activity, including trypsin, contributes to epithelial barrier dysfunction and altered microbial ecology^8,47^. Moreover, members of the gut microbiota actively modulate disease-associated trypsin activity through extracellular proteolytic mechanisms^46,48^, suggesting that adaptation to proteolytic environments is an important determinant of bacterial fitness.

To test this hypothesis, we exposed wild-type *P. vulgatus* and the isogenic *Δdpp11a* mutant to increasing concentrations of serine proteases i) porcine trypsin and ii) proteinase K in minimal media. Both proteases inhibited growth in a dose-dependent manner (**Fig. S4a,b**). Dose-response modelling indicated that the trypsin IC20 (0.08 mg mL⁻¹) approximated the lower range of trypsin-equivalent protease activity reported in healthy human feces, whereas IC50-IC70 values (0.3-0.7 mg mL⁻¹) overlapped with the elevated protease activities reported in IBD and UC ^15,46^. Across both low (approximately IC20-IC40) and high (approximately IC50-IC80) protease concentrations, loss of Dpp11a modestly but significantly increased susceptibility to both trypsin (〜10% increased growth inhibition of *Δdpp11a* only at high concentrations, p =0.02: two-sided Wilcoxon rank-sum test. **Fig. 4a–b**) and proteinase K (〜15% increased growth inhibition of *Δdpp11a* at both low and high concentrations: p = 0.018 and p = 0.04, respectively, two-sided Wilcoxon rank-sum test. **Fig. S4c,d**). Thus, Dpp11a partially protects *P. vulgatus* from diverse extracellular serine proteases and may facilitate adaptation to proteolytic environments such as the inflamed intestine. In contrast, deletion of *dpp11a* did not increase susceptibility to the host antimicrobial effectors lysozyme, LL-37, hBD2, or hBD3, nor did it impair persistence within a defined microbial community (**Fig. S5a,b**), indicating that Dpp11a does not function in general stress resistance. Together, these findings indicate that Dpp11a confers a modest but specific level of protection against extracellular serine proteases, consistent with a role in promoting *P. vulgatus* resilience under protease-rich conditions such as those encountered during intestinal inflammation.

**Figure 4.**
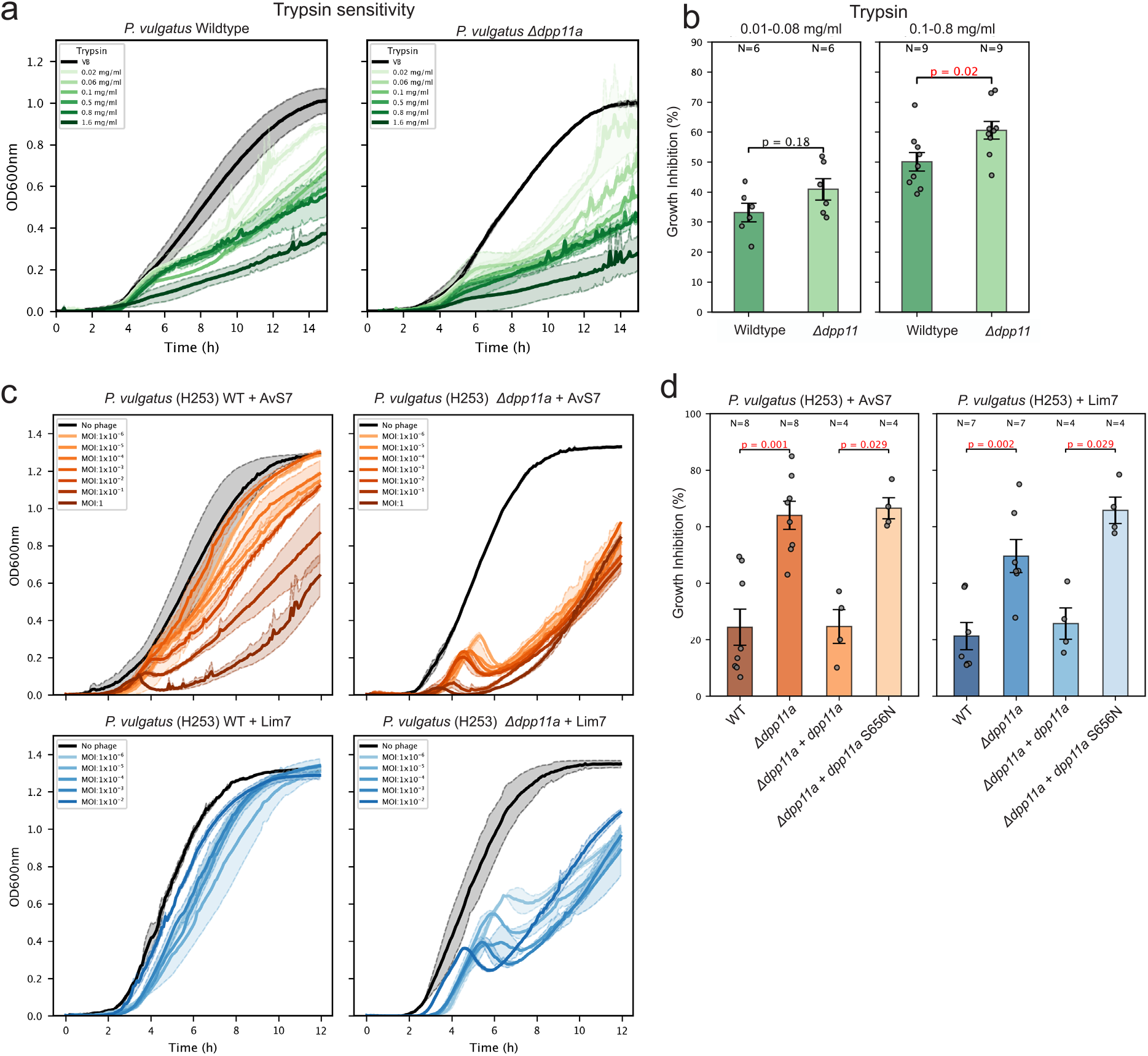
Dpp11a protects *P. vulgatus* against extracellular trypsin and bacteriophage infection. **a)** Representative growth curves of wild-type *P. vulgatus*^ATCC 8482^ and an isogenic Δ*dpp11a* mutant cultured in the presence of increasing concentrations of porcine trypsin (0.02-1.6 mg ml⁻¹) or untreated control (black). Technical replicates are shown as thin coloured lines and the mean as a thick line; shaded regions indicate ± s.e.m. Representative of three independent experiments (replicates in Supplementary Data). **b**) Growth inhibition of wild-type and Δ*dpp11a* strains at low (0.01-0.08 mg ml⁻¹) and high (0.1-0.8 mg ml⁻¹) trypsin concentrations. Growth inhibition (%) was calculated from the AUC relative to untreated controls. Bars represent mean ± s.e.m.; points represent individual measurements pooled from at least three independent experiments. N denotes the number of data points per group, combined from three independent experiments with varying numbers of concentrations tested per experiment. A two-sided Mann-Whitney U test was used to test significance. **c**) Representative growth curves of wild-type *P. vulgatus*^H253^ and an isogenic Δ*dpp11a* mutant following infection with AvS7 (upper) or Lim7 (lower) bacteriophages across the indicated multiplicities of infection (MOIs); uninfected controls are shown in black. Technical replicates are shown as thin coloured lines and the mean as a thick line; shaded regions indicate ± s.e.m. Representative of three independent experiments (replicates in Supplementary Data). **d**) Growth inhibition following infection with AvS7 (left) or Lim7 (right) at a low MOI (10⁻^4^) in wild-type *P. vulgatus* ^H253^, the Δ*dpp11a* mutant, a chromosomally complemented Δ*dpp11a* strain (Δ*dpp11a*::pNBU2-*dpp11a*), and a complemented strain expressing a catalytically inactive Dpp11a variant (Δ*dpp11a*::pNBU2-*dpp11a*^S656N^). Growth inhibition was calculated from the AUC relative to uninfected controls at 12 h. Bars represent mean ± s.e.m.; data points represent individual biological replicates binned according to the indicated concentration ranges tested from at least three independent experiments. *N* denotes the number of data points per group, combined from three independent experiments with varying numbers of concentrations tested per experiment. A two-sided Mann-Whitney U test was used to test significance.

IBD is additionally characterized by expansion of intestinal bacteriophages, particularly members of the *Caudovirales* order, which coincide with reductions in *Bacteroidia* abundance and altered gut microbial ecology^16,19,20^. Since extracellular proteases have previously been implicated in anti-phage defense systems^49,50^, we next asked whether Dpp11a also contributes to resistance against bacteriophage infection. We therefore challenged wild-type and Δ*dpp11a P. vulgatus* CLA-AA-H253, a phage-permissive isolate, with two lytic Siphoviridae phages, AvS7 and Lim7. Across a range of multiplicities of infection (MOIs), *dpp11a* deletion led to increased sensitivity to predation by both phages, with the mutant exhibiting markedly impaired growth relative to the wild-type (**Fig. 4c**). To confirm this phenotype is specifically attributable to Dpp11a, we generated strains complemented with either wild-type *dpp11a* or a catalytically inactive *dpp11a*^S656N^ allele. TAMRA-FP labeling verified restoration of Dpp11a activity only in the strain complemented with wild-type *dpp11a* (**Fig. S6a**). Representative plaque assays, the complete phage genomes, and infection growth curves for the complemented strains are shown in **Fig. S6b–d**. Consistent with the growth inhibition observed in the *dpp11a* deletion mutant (〜 two-fold increase in Growth Inhibition for both AvS7 p = 0.001, two-sided Mann-Whitney U test and Lim7 p = 0.002, two-sided Mann-Whitney U test), complementation with wild-type *dpp11a*, but not the catalytically inactive *dpp11a*^S656N^ allele, restored resistance to both phages (**Fig. 4d**. AvS7 p = 0.029, two-sided Mann-Whitney U test. Lim7 p = 0.029, two-sided Mann-Whitney U test). Together, these findings identify Dpp11a as a catalytically active anti-phage defense factor, revealing an unexpected role for a periplasmic/extracellular serine protease in protecting *P. vulgatus* from bacteriophage infection.

### UC-associated transcriptional changes implicate Dpp11a in phage-dependent epithelial barrier modulation

Given the elevated protease activity and bacteriophage blooms associated with UC, we next asked whether the Dpp11a defense pathway is altered during human disease. Previous studies have reported expansion of intestinal tailed bacteriophages together with disruption of *Bacteroidales* populations in both UC and Crohn’s disease^16,19,20^. Analysis of metatranscriptomic data from the Lloyd-Price *et al*.^42^ IBD cohort revealed significantly increased expression of both *P. vulgatus dpp11a* (p = 0.045, two-side Mann-Whitney U test, **Fig. 5a)** and the downstream gene *dpp11b* in UC compared with non-IBD controls (p = 0.023, two-side Mann-Whitney U test, **Fig. S7a**). In contrast, other DPP family members, including *dpp4a*, *dpp4b*, and *dpp7*, showed little or no disease-associated change (**Fig. S7a**). The concordant regulation of *dpp11a* and *dpp11b* is consistent with possible coordinated expression of the Dpp11 locus, potentially reflecting their organization within the same operon, although direct co-transcription remains to be established. Transcripts derived from combined *P. vulgatus*-associated bacteriophages sequences were also significantly elevated in UC relative to non-IBD controls (p = 2.0×10^-8^, two-side Mann-Whitney U test, **Fig. 5b**), indicating that increased expression of Dpp11a coincides with enhanced phage activity in the inflamed gut.

**Figure 5.**
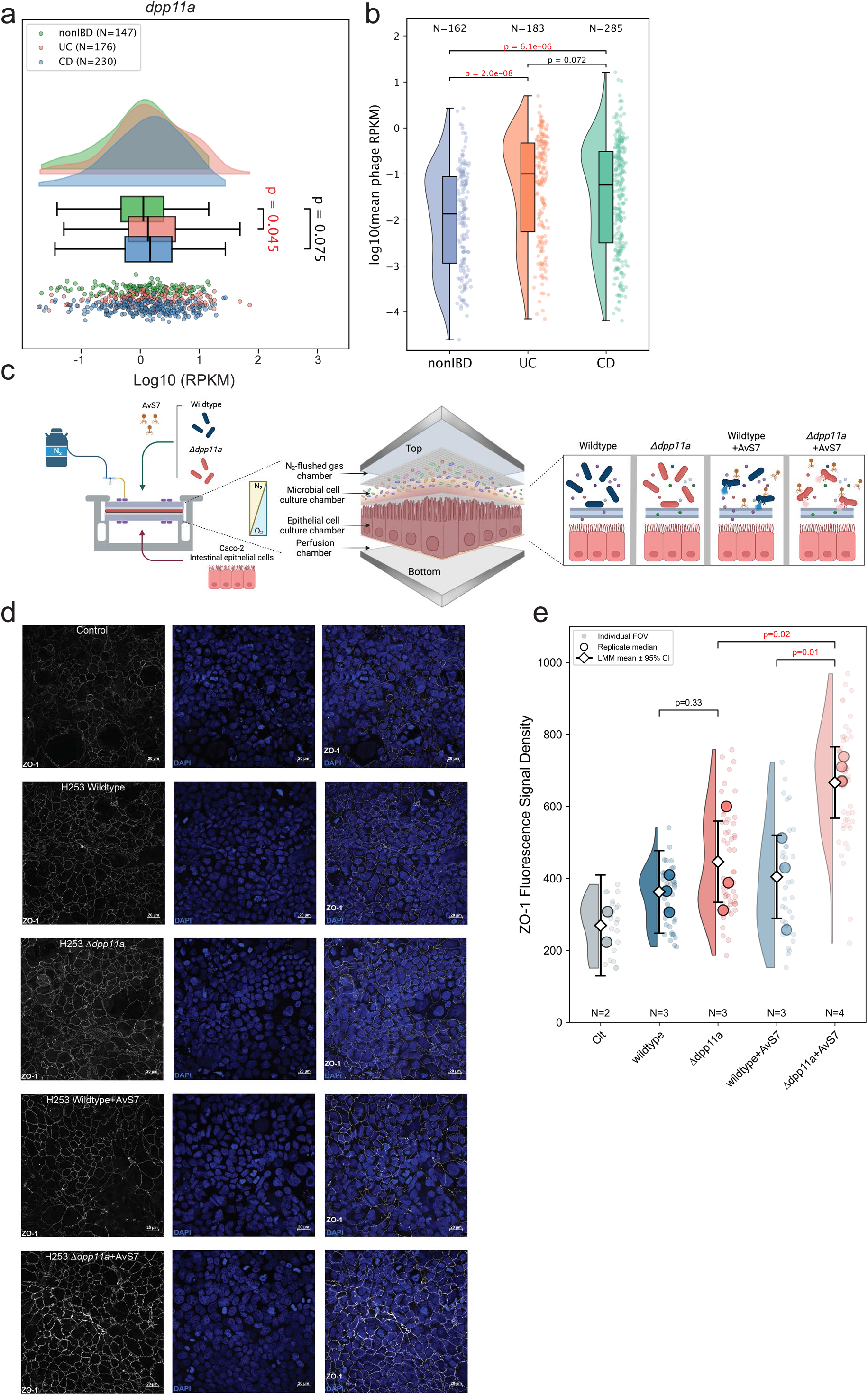
Altered Dpp11a and phage transcripts in UC are linked to Dpp11a-dependent modulation of epithelial barrier integrity upon phage infection. **a**) Expression of *P. vulgatus dpp11a* in metatranscriptomic datasets from healthy non-IBD controls, patients with ulcerative colitis (UC), and patients with Crohn’s disease (CD)^42^. Density plots show the distribution of transcript abundance (log10 RPKM). Boxes indicate the interquartile range (IQR) with the center line denoting the median, whiskers extend to the most extreme values within 1.5 × IQR, and points represent individual samples. Statistical significance was assessed using two-sided Mann-Whitney U tests. **b**) Metatranscriptomic analysis of the same cohort showing pooled transcript abundance of *P. vulgatus*-associated *Caudovirales* bacteriophages (log10 mean phage RPKM). Raincloud plots showing the distribution of log10(mean phage RPKM) across nonIBD, UC and CD samples. Half-Violin plots show the data distribution; boxes indicate the IQR with the center line denoting the median, whiskers extend to the most extreme values within 1.5 × IQR, and points represent individual samples. Sample sizes are indicated above each group. Statistical significance was assessed using two-sided Mann-Whitney U tests. **c**) Schematic of the HuMiX microfluidic gut-on-a-chip system used to investigate bacterium-phage-host interactions. *P. vulgatus* ^H253^ strains and bacteriophage AvS7 were introduced into the microbial chamber, while differentiated Caco-2 intestinal epithelial cells were cultured in the adjacent epithelial chamber under continuous perfusion and physiologically relevant oxygen gradients. **d**) Representative maximum-intensity projections of confocal Z-stacks showing Caco-2 epithelial monolayers stained for the tight-junction protein ZO-1 (white) and nuclei (DAPI, blue) following exposure to the indicated bacterial and phage treatments. Scale bars, 20 μm. Images are representative of three independent experiments. **e**) Quantification of ZO-1 fluorescence density from the experiments shown in **d**. ZO-1 fluorescence was normalized to the segmented DAPI-positive nuclear area within each field of view (FOV). Light data points represent individual FOVs, large circles denote biological replicate medians, and diamonds indicate model-estimated marginal means ± 95% Wald confidence intervals from a Gaussian linear mixed-effects model. Experimental condition was included as a fixed effect and biological replicate (10-25 FOV’s per replicate) as a random intercept. Pairwise comparisons were performed using the Satterthwaite approximation for denominator degrees of freedom. Exact *P* values are shown. *N* indicates the number of independent biological replicates.

Because bacteriophages are increasingly recognized as modulators of epithelial barrier function and inflammatory responses^51–56^, we next investigated whether Dpp11a-dependent responses to phage infection influence host epithelial physiology using the HuMiX microfluidic gut-on-a-chip system^57^ (**Fig. 5c**). We focused on epithelial tight-junction organization as a readout for barrier-associated responses, using ZO-1 fluorescence to assess changes in junctional organization. Co-cultivation with wild-type or Δ*dpp11a P. vulgatus* in the absence of phage produced comparable epithelial morphology, nuclear shape, ZO-1 fluorescence, cell viability, and bacterial abundance, indicating that loss of Dpp11a does not measurably alter bacterial colonization or epithelial homeostasis under basal conditions (**Fig. 5d**,**e**; **Fig. S7b–d**). In contrast, phage infection elicited genotype-dependent epithelial responses. AvS7 challenge of the Δ*dpp11a* mutant resulted in significantly greater epithelial ZO-1 fluorescence than phage infection of wild-type bacteria (Gaussian linear mixed-effects model with Satterthwaite-adjusted pairwise comparisons; p = 0.012 **Fig. 5d-e**), consistent with altered tight-junction organization. This difference occurred without detectable changes in epithelial nuclear morphology, cell viability, or bacterial abundance (**Fig. S7b–d**), arguing against altered colonization or cytotoxicity as the primary explanation. These results link a disease-associated bacterial phage-defence pathway with epithelial responses during phage infection, revealing consequences of Dpp11a activity that extend beyond bacteria–phage interactions to the host interface.

## Discussion

Microbiome-derived proteases are increasingly recognized as regulators of host physiology, yet the ecological functions remain poorly understood. Here, we identify Dpp11a as the dominant active serine protease produced by *P. vulgatus* and show that it protects against extracellular proteolytic stress and bacteriophage infection. We further show that bacterial responses to phage challenge influence epithelial junctional remodeling. Together, these findings expand the functional repertoire of bacterial DPP’s and identify microbial phage defense pathways as an underappreciated mechanism through which intestinal ecological interactions within the gut microbiome can impact host physiology.

Dipeptidyl peptidases encoded by *Bacteroidales* have largely been studied as nutrient acquisition enzymes, particularly in *Porphyromonas gingivalis*, wherein Dpp11 contributes to amino acid utilization and bacterial growth^29^. In contrast, our findings suggest functional specialization between the two closely related Dpp11 homologues encoded by *P. vulgatus*. Whereas Dpp11b promoted growth under nutrient-limiting conditions, Dpp11a was dispensable for growth, yet protected against extracellular proteases and bacteriophage infection. The adjacent genomic organization of *dpp11b* and *dpp11a*, together with their sequence and functional divergence, is consistent with a gene duplication event followed by neofunctionalization. We therefore speculate that Dpp11a represents an ancestral metabolic peptidase that has been evolutionarily repurposed into a factor aiding in bacterial defense.

The dual requirement for Dpp11a catalytic activity in extracellular protease tolerance and phage resistance raises the intriguing possibility that these phenotypes share a common mechanistic basis. One possibility is that Dpp11a directly processes extracellular phage proteins, alternatively it may modify bacterial substrate proteins that enhance envelope resilience or alter phage receptor accessibility during infection. The complex and dynamic ecological context of the intestine provides a plausible explanation for the evolution of this function. During inflammatory bowel disease, luminal trypsin activity is elevated and gut bacteria can actively regulate intestinal protease abundance^46^, exposing abundant *Bacteroidales* to sustained proteolytic stress while simultaneously experiencing increased phage predation. Such combined selective pressures may have favored the recruitment of a catabolic protein/peptide hydrolase into bacterial defense. This evolutionary strategy is increasingly recognized across bacterial immunity, where pre-existing cellular machinery have been co-opted for antiviral defence. ^58,59^.

The coexistence of elevated protease activity and bacteriophage blooms is a defining feature of UC. Consistent with this environment, we observed increased expression of both *dpp11a* and *P. vulgatus*-associated phage transcripts in meta-transcriptomes from the Lloyd-Price IBDMDB cohort^42^, suggesting activation of this defense pathway during disease. Although *P. vulgatus* abundance varies between IBD cohorts, increasing evidence indicates that strain-specific functional traits are more informative than taxonomic abundance alone ^13^. Our findings therefore suggest that differences in phage susceptibility may represent an additional layer through which the microbiome influences disease. Consistent with this idea, phage infection elicited distinct epithelial responses depending on bacterial genotype despite comparable bacterial abundance and epithelial viability. Increased ZO-1 remodelling following infection of the phage-sensitive Δ*dpp11a* mutant raises the possibility that enhanced phage susceptibility alters the release or presentation of bacterial or phage-derived products through phage-induced lysis, which appear to be subsequently sensed by the epithelium. Although the underlying mechanism and signals remain unknown, these findings suggest that host tissues likely respond not only to bacteria or phages individually, but also indirectly to bacterial defense pathways that determine the outcome of bacterial-phage-host interactions.

Several limitations of our study should be considered. Our epithelial co-culture experiments cannot distinguish whether the observed changes in epithelial remodeling arise from Dpp11a itself, phage-induced release of bacterial products, or secondary alterations in bacterial physiology. Likewise, the molecular substrates responsible for both protease tolerance and phage resistance remain unknown. Future studies combining substrate identification, comparative genomics, and *in vivo* models will be important for determining how broadly Dpp11-mediated defense is conserved across *Bacteroidales* and whether related pathways contribute to intestinal disease.

Together, our findings identify Dpp11a as a multifunctional bacterial defence factor that contributes to resistance against both extracellular proteases and bacteriophage infection. More broadly, they position bacterial phage defence as an underappreciated nexus between ecological pressures in the gut and host epithelial responses. This framework expands our view of host-microbiome interactions beyond microbial composition and metabolic cross-talk, highlighting phage susceptibility as a potential determinant of epithelial responses to gut microbiota.

## Author Contributions

Conceptualization, N.Q. and J.S.; Methodology, N.Q., M.S.B., K.B., M.G., L.M., C.D.R., C.P., P.W., E.M.B., and J.S.; Investigation, N.Q., M.S.B., K.B., M.G., L.M., C.P., N.T., M.S.M. A.K. T.C.A.H., B.M. and E.M.B.; Formal Analysis, N.Q., M.S.B., T.C.A.H. M. S. M. and E.M.B.; Resources, B.S., C.G.P.V., A.T., and B. O. S., P.W., T.C.; Data Curation, N.Q.; Writing Original Draft, N.Q. and J.S.; Writing, Reviewing & Editing, all authors; Visualization, N.Q.; Supervision, J.S.; Funding Acquisition, N.Q., T.C. and J.S.

## Resource Availability Lead Contact

Further information and requests for resources and reagents should be directed to and will be fulfilled by the Lead Contact, Joel Selkrig.

## Material availability

Bacterial strains and plasmids generated in this study are available from the Lead Contact upon reasonable request.

## Data and Code Availability

Mass spectrometry proteomics data generated in this study have been deposited to the ProteomeXchange Consortium via the PRIDE partner repository under accession number PXD081763 and will be made publicly available as of the date of publication.

Original code used for data analysis is available from the Lead Contact upon reasonable request prior to publication.

Any additional information required to reanalyze the data reported in this manuscript is available from the Lead Contact upon request.

## Acknowledgments

We thank all members of the Selkrig lab for helpful discussions throughout the project. This research project was partly funded by the START-Program of the Faculty of Medicine RWTH Aachen University (awarded to J.S. and N.Q.) and by the German Research Foundation (DFG, Deutsche Forschungsgemeinschaft) Project-ID 564421876 awarded to J.S.. The co-culture part of this work was supported by JUMP grant (19303451/HuMiX-HT) all funded by the Luxembourg National Research Fund and awarded to P.W. B.S. received funding by the European Research Council (ERC) under the European Union’s Horizon 2020 research and innovation program (no. 865615), the German Research Foundation (DFG, German Research Foundation, SFB 1371, no. 395357507), and the German Center for Infection Research (DZIF). BOS acknowledges funding from the Swedish Research Council (Vetenskapsrådet; #2021-06602). We thank Dimitris Kapsokalyvas for valuable advice on microscopy image quantification. Isolation of strains used in this study were supported by SFB1382 Gut-liver axis, DFG no. 424795268, project Q02, SFB1371 Microbiome signatures, DFG no. 395357507, project Z01 awarded to T. C. We thank members of the IZKF Proteomics Core Facility at RWTH University Hospital Aachen for technical support.

## Materials and Methods

### Bacterial strains and growth conditions

Bacterial strains are listed in **Table S4**. *Phocaeicola vulgatus^ATCC^* ^8482^ was used as the reference strain for most experiments, whereas *P. vulgatus^CLA-AA-H^*^253^ (*P. vulgatus^H^*^253^, H253) and derived isogenic mutants were used for phage infection assays. *P. vulgatus* strains were cultured anaerobically at 37°C in modified Gifu Anaerobic Medium (mGAM; HyServe GmbH & Co. KG/Nissui Pharmaceuticals, cat. no. 1005433-001). Protease sensitivity assays were performed in Varel and Bryant (VB) minimal medium^60^ to avoid interference from peptides present in rich medium. Media were pre-reduced for at least 24 h before use in an anaerobic chamber containing 2% H2, 12% CO2, and balance N2. *E. coli* DATC ^61^ used for cloning was grown aerobically in LB medium supplemented with 0.3 mM diaminopimelic acid at 37°C with shaking. Plasmids derived from pNBU2 or pLGB13 were maintained with 100 μg/mL ampicillin. Bacterial stocks were stored at −80°C in 15% glycerol.

### Molecular cloning and strain construction

Genetic manipulation of *P. vulgatus* was performed as previously described with minor modifications ^62^. Deletion of *dpp11a*(*BVU_2253*) in *P. vulgatus^ATCC^* ^8482^ and *P. vulgatus^H^*^253^ was performed by pLGB13-mediated allelic exchange. Approximately 1 kb upstream and downstream homology arms flanking *dpp11a* were amplified from *P. vulgatus^ATCC^* ^8482^ genomic DNA using primer pairs JSOL_0123/JSOL_0124 and JSOL_0125/JSOL_0126, respectively, and assembled into PCR-linearized pLGB13 generated with primers JSOL_0121/JSOL_0122 (all primers listed in **Table S5)**. For complementation and overexpression, wildtype *dpp11a*, catalytically inactive *dpp11a ^S^*^656^*^A^*, C-terminally 2xHA-tagged *dpp11a,* and C-terminally 3xFLAG-tagged *dpp11a* were cloned downstream of the constitutive P1E6 promoter in a pNBU2-based integration vector. Untagged constructs were generated using pNBU2 linearized with primers JSOL_0013/JSOL_0014, whereas the 2xHA- and 3xFLAG-tagged constructs were generated using tag-containing pNBU2 backbones linearized with JSOL_0198/JSOL_0014 and JSOL_0200/JSOL_0014, respectively. The S656N catalytic-site substitution was introduced by overlap-extension PCR using primer pairs JSOL_0084/JSOL_0179 and JSOL_0178/JSOL_0085. Epitope tags were fused in frame immediately upstream of the native *dpp11a*stop codon. All constructs were assembled using Gibson Assembly (New England Biolabs, E2621L) according to the manufacturer’s instructions, propagated in *E. coli* DATC under ampicillin selection, and verified by colony PCR and Sanger sequencing. Primer sequences, mutagenesis strategy, and PCR conditions are provided in **Tables S5 and S6.**

pLGB13- and pNBU2-derived plasmids^62^ were conjugated into *P. vulgatus* using the *E. coli* DATC. Transconjugants were selected anaerobically on mGAM agar containing 50 µg/mL erythromycin. pNBU2 integrants were retained as erythromycin-resistant clones. For pLGB13-mediated allelic exchange, erythromycin-resistant merodiploids were passaged overnight in liquid mGAM without antibiotics and subsequently counter-selected on mGAM agar supplemented with 100 ng/mL anhydrotetracycline to promote plasmid resolution. Erythromycin-sensitive colonies were screened by PCR, and deletion of *dpp11a* was confirmed using primers located outside the recombination arms. Screening and confirmation primers are listed in **Tables S5 and S6**.

### Activity-based protein profiling and proteomics

#### TAMRA-FP labeling

Active serine hydrolases were detected using the ActivX TAMRA-FP serine hydrolase probe (TAMRA-FP; Thermo Fisher Scientific, cat. no. 88318). *P. vulgatus^ATCC^* ^8482^ wildtype and the isogenic *Δdpp11a* mutant were grown anaerobically in mGAM. For whole-cell ABPP analysis, 1 mL of overnight culture was harvested by centrifugation at 8,000 x *g* for 5 min at room temperature. Cell pellets were washed twice with 1 mL PBS and resuspended in an equal volume of PBS. For cell pellet versus supernatant fractionation and OMV analysis, late-exponential-phase cultures were used to minimize stationary-phase autolysis. Cultures were centrifuged to separate cell pellets and supernatants. Supernatants were sterile-filtered through 0.2 μm membranes (SFCA membrane; CORNING®, cat. no. 431219), and cell pellets were washed and resuspended in PBS. For OMV analysis, filtered supernatants were concentrated using 3 kDa MWCO (Amicon® Ultra, Merck Millipore, cat. no. UFC900324) centrifugal filters and subsequently ultracentrifuged at 100,000 x *g*for 1 h at 4°C to separate OMV-enriched pellets from soluble supernatant fractions.

Samples were incubated with 1 μM TAMRA-FP for 30 min at room temperature in the dark. Reactions were stopped by addition of Laemmli loading buffer ^63^, followed by heating at 95°C for 10 min. Labeled proteins were resolved by 10% SDS-PAGE and visualized by in-gel fluorescence scanning on a ChemiDoc XRS+ system (Bio-Rad) using a Cy3-compatible filter suitable for TAMRA fluorescence detection, with excitation and emission maxima of approximately 552 nm and 575 nm, respectively. Protein loading was assessed by anti-GroEL immunoblotting. After fluorescence imaging, proteins were transferred to 0.45 μm nitrocellulose membranes (Merck Millipore, cat. no. GE10600016) using a semi-dry blotting system. Membranes were blocked with 5% skim milk in TBST for 1 h at room temperature and incubated overnight at 4°C with anti-GroEL antibody (Sigma-Aldrich, cat. no. GG6532; 1:80,000). After washing, membranes were incubated with HRP-conjugated anti-rabbit secondary antibody (Abcam, cat. no. ab6734; 1:20,000) for 1 h at room temperature. Signals were detected using ECL substrate (Bio-Rad, cat. no. 1705060) and imaged with a ChemiDoc XRS+ system (Bio-Rad).

### PMSF competition assay

To validate TAMRA-FP labeling specificity, pellets harvested from overnight culture were pre-incubated with 1 mM phenylmethylsulfonyl fluoride (PMSF) or solvent control for 30 min on ice before TAMRA-FP labeling. Samples were then labeled with 1 μM TAMRA-FP and analyzed by in-gel fluorescence as described above.

### Competitive TAMRA-FP and Biotin-FP labeling

Competitive ABPP was performed using TAMRA-FP and ActivX Desthiobiotin-FP serine hydrolase probe (Biotin-FP; Thermo Fisher Scientific, cat. no. 88317). *P. vulgatus^ATCC^* ^8482^ cultures were grown to mid-log phase, and cell pellets and concentrated culture supernatants were prepared as described above. Samples were pre-incubated with 1 μM TAMRA-FP or PBS for 30 min at room temperature in the dark, followed by labeling with 1 μM Biotin-FP for 30 min. For western blot detection, proteins were denatured in loading buffer, excess Biotin-FP was removed using 7 kDa MWCO Zeba™ spin desalting columns (Thermo Fisher Scientific, cat. no. 89877) according to the manufacturer’s instructions, and biotin-labeled proteins were detected using HRP-conjugated streptavidin (Cell Signaling, cat. no. #3999; 1:2000 dilution in 5% BSA/TBST). BSA was used for membrane blocking and antibody dilution to minimize background from endogenous biotin in milk.

### Affinity enrichment of Biotin-FP-labeled proteins

For enrichment of Biotin-FP-labeled proteins, denatured samples were desalted and diluted to 0.2% SDS in PBS. Biotin-labeled proteins were captured using NeutrAvidin-agarose beads (50 µL per sample; Thermo Fisher Scientific, cat. no. 29200) for 1.5 h at room temperature with end-over-end rotation. Beads were washed with 0.2% SDS-containing PBS to remove non-specifically bound proteins and processed for on-bead digestion.

### On-bead digestion and peptide cleanup

NeutrAvidin-agarose beads were washed with 2x PBS followed by 50 mM TEAB (2,000 x *g*, 1 min, room temperature). Beads were resuspended in 25 mM TEAB containing 10 mM TCEP and 40 mM chloroacetamide and incubated for 30 min at 55°C for reduction and alkylation. After washing with 50 mM TEAB, proteins were digested on beads in 50 mM TEAB containing MS-grade trypsin (Thermo Fisher Scientific, cat. no. 13464189) for 14 h at 37°C. The digestion mixture was centrifuged, and the peptide-containing supernatant was transferred to a fresh tube and acidified with formic acid to 0.5%. Peptides were desalted on C18 StageTips, washed with 0.1% formic acid, eluted twice with 0.1% formic acid in 80% acetonitrile, and dried under vacuum.

### TMT labeling

Peptides were labeled with TMTpro reagents using an optimized protocol based on Zecha *et al.*^55^. Peptides were resuspended in 50 mM HEPES (pH 8), and labeling was performed in small reaction volumes with a final TMT concentration of 11.8 mM and a final acetonitrile concentration of 20%. Reactions were incubated for 1 h at room temperature, quenched with hydroxylamine, pooled, desalted using Oasis HLB μElution plates, eluted with 60% methanol/1% formic acid, and dried under vacuum.

### LC-MS/MS analysis and data processing

Peptides were resuspended in 3% formic acid/1% acetonitrile and analyzed by LC-MS/MS using an UltiMate 3000 RSLCnano system (Thermo Fisher Scientific) coupled to an Orbitrap Exploris 480 mass spectrometer meter (Thermo Fisher Scientific, cat. no. BRE725533). Peptides were separated on a C18 analytical column using a 100 min acetonitrile gradient and analyzed in positive ion mode using data-dependent acquisition (DDA). Full MS scans were acquired over 375-1,500 m/z at 120,000 resolutions (AGC=300%), followed by MS/MS acquisition of peptides with charge states 2-5.

Raw files were converted to mzML and processed with MSFragger (v3.8) against the *P. vulgatus^ATCC^* ^8482^ UniProt proteome (UP000002861), supplemented with streptavidin (UniprotID: P22629) and common contaminants and reverse decoy sequences. Protein identification and quantification required at least one unique peptide, and peptide-level FDR was controlled at 1%.

Downstream analysis was performed in R (v2021.09.2) using the limma (v3.54.2) and tidyverse packages including dplyr (v1.1.1) and ggplot2 (v3.4.2). Reporter ion intensities were normalized by total sample loading and log2-transformed before statistical analysis. Differential abundance was assessed using linear models implemented in limma with moderated t tests and empirical Bayes variance estimation. p values were adjusted for multiple testing using the Benjamini-Hochberg method.

*P. vulgatus^ATCC^* ^8482^ protein abundance analysis

For protein abundance analysis, *P. vulgatus^ATCC^* ^8482^ was grown anaerobically in mGAM to mid-log phase. Cells were harvested by centrifugation at 6,000 x *g* for 10 min at 4°C, washed with 50 mM HEPES (pH 8.0), and lysed in 50 mM HEPES containing 1% SDS by heating at 95°C for 10 min.

### Sample preparation

Samples were lysed in 4x lysis buffer (50 mM HEPES [pH 8], 4% SDS, 160 mM chloroacetamide, and 40 mM TCEP) for 5 min at 95°C. Nucleic acids were degraded with Benzonase nuclease (0.5 units per μg sample) for 30 min at 37°C. Proteins were captured on SP3 beads (1:1 hydrophobic/hydrophilic carboxyl-coated Sera-Mag Speed Beads; GE Healthcare, cat. nos. 45152105050250 and 65152105050250) using 50% acetonitrile, washed with 80% ethanol, and digested on beads in 50 mM TEAB containing trypsin (Serva) and LysC (Wako) for 14 h at 37°C. Peptide-containing supernatants were collected and dried under vacuum.

### LC-MS/MS analysis and data processing

Peptides were resuspended and analyzed as above using a 160 min gradient. Full MS scans were acquired over 400-1,600 m/z at 120,000 resolution (AGC = 100%), followed by MS/MS acquisition of peptides with charge states 2-5. Raw files were converted to mzML and processed with MSFragger (v3.8) against the *P. vulgatus^ATCC^* ^8482^ UniProt proteome (UP000002861), Bos taurus (UP000009136), and Sus scrofa (UP000008227) UniProt databases with contaminants and reverse sequences. Protein identification and quantification required at least one unique peptide, and peptide-level FDR was controlled at 1%. Downstream analysis was performed in R as above. Trimmed Mean of M-values (TMM) normalization was applied to account for composition biases. TMM normalization factors were calculated using the calcNormFactors function from the edgeR Bioconductor package^64^. Differential abundance was assessed using linear models implemented in limma as described above.

### Osmotic shock fractionation

Subcellular localization of Dpp11a was assessed by osmotic shock fractionation as previously described ^65–67^ using a *P. vulgatus^ATCC^* ^8482^ *Δdpp11a* strain expressing chromosomally integrated Dpp11a-2xHA. Cells were grown anaerobically in mGAM to mid-log phase and harvested at 8,000 x *g* for 10 min at 4°C. Pellets were resuspended in ice-cold hypertonic buffer containing 20 mM Tris-HCl, pH 7.5, 20% sucrose, and 1 mM EDTA and incubated on ice for 15 min. Cells were then pelleted, gently resuspended in ice-cold hypotonic buffer containing 20 mM Tris-HCl, pH 7.5, and 1 mM EDTA, and incubated on ice for 20 min. After centrifugation at 12,000 x *g* for 15 min at 4°C, the supernatant was collected as the osmotic shock fraction, and the pellet was retained as the remaining cell-associated fraction.

The osmotic shock supernatant was further ultracentrifuged at 100,000 x *g* for 1 h at 4°C to separate soluble periplasmic proteins from membrane-associated components. The resulting supernatant and pellet were collected as soluble and membrane-enriched fractions, respectively. Fractions were analyzed by 10% SDS-PAGE and western blotting. Dpp11a-2xHA was detected using anti-HA antibody (BioLegend, cat. no. 902301; 1:1,000), and GroEL was detected as a cytoplasmic marker using anti-GroEL antibody as described above.

### Immunogold transmission electron microscopy

*P. vulgatus^ATCC^* ^8482^ *Δdpp11a* strain expressing C-terminally 3xFLAG-tagged Dpp11a was grown anaerobically overnight in mGAM at 37°C. Cell pellets were harvested by centrifugation at 8,000 x *g* for 5 min, the supernatant was removed, and pellets were fixed in 4% paraformaldehyde. Fixed cells were embedded in Lowicryl HM20 (Polysciences, Warrington, PA, USA) via freeze substitution after high-pressure freezing. Freeze substitution was performed with methanol dehydration steps at -90 °C and ethanol dehydration steps at -50 °C, down to 0 °C. Polymerization was done at -50 °C via UV light.

Immune labeling was performed on ultrathin sections. Sections were incubated with 50 mM glycine for 20 min to quench aldehyde residues, blocked with 5% BSA and incubated with anti-FLAG-Tag-antibody (Merck, cat. co. F1804) 1:10 diluted in 1% BSA/PBS overnight at 4° C. After washing with 0.05% Tween-20 in PBS, sections were incubated for 1 h with 12 nm gold-conjugated anti-mouse IgG secondary antibody (Jackson ImmunoResearch Europe Ltd., cat. no. 715-205-150; 1:40) diluted in 1% BSA/PBS.

Contrast was enhanced by staining with 0.5% uranyl acetate and 1% lead citrate (both EMS, Munich, Germany). Samples were examined using a Hitachi HT7800 transmission electron microscope (Hitachi, Tokyo, Japan) operating at an acceleration voltage of 100 kV.

### Defined community dynamics assay

A defined microbial community (DMC) comprising nine bacterial strains (listed in **S4**) isolated from non-IBD donors ^68^ was supplemented with either *P. vulgatus ^ATCC^* ^8482^ wild-type or the isogenic *Δdpp11a* mutant. Equal volumes of overnight cultures were combined to assemble the community, aliquoted in 15% glycerol, and stored at −80°C. For each independent experiment, DMC stocks were inoculated 1:10 into fresh mGAM medium and cultured anaerobically at 37°C. Communities were passaged daily (1:10 dilution) for five consecutive days, and samples were collected every 24 h, pelleted by centrifugation, and stored at −80°C for downstream analysis.

### 16S rRNA gene amplicon sequencing and analysis

Metagenomic DNA was extracted using a modified protocol based on Godon *et al.* ^69^, incorporating mechanical bead beating and column-based purification. The V3-V4 region of the 16S rRNA gene was amplified and sequenced on an Illumina MiSeq platform (2 × 300 bp) as previously described ^70–72^. Raw reads were processed using the IMNGS platform based on the UPARSE pipeline ^73^, and operational taxonomic units were taxonomically assigned.

Relative abundances were calculated following Rhea normalization ^72^. OTUs were aggregated at the lowest assigned taxonomic level. Community composition retained between consecutive sampling time points was calculated as the sum of the minimum relative abundance of each taxon shared between the two time points (i.e., Σ min[relative abundance at time 1, relative abundance at time 2]), expressed as a percentage of the community. Downstream analyses and data visualization were performed in Python (v3.9).

### Protease sensitivity assay

Protease sensitivity of *P. vulgatus^ATCC^* ^8482^ wildtype and the isogenic *Δdpp11a* mutant was assessed using porcine trypsin (Sigma-Aldrich, cat. no. T5266) and proteinase K (PanReac AppliChen, cat. no. A3830,0100). Strains were grown anaerobically overnight in mGAM, diluted to an initial OD600nm of 0.01 in Varel and Bryant minimal medium ^60^, and dispensed into U-bottom 96-well plates containing the indicated concentrations of porcine trypsin (Sigma-Aldrich, cat. no. T5266), or proteinase K (PanReac AppliChen, cat. no. A3830,0100). Plates were sealed with gas-permeable membranes (Sigma-Aldrich, Breathe-Easy®, cat. no. Z380059) and incubated anaerobically at 37°C with continuous shaking at 350 rpm. OD600nm was measured every 3 min for 24 h using an Alto Cerillo plate reader.

For each biological replicate, OD600nm values from two technical replicate well were averaged to generate one mean growth curve per condition per independent experiment. Growth curves were baseline-normalized by subtracting the initial OD600nm value, and the area under the curve (AUC) was calculated by numerical integration using custom code in Python (v3.9). The integration window was defined based on the time at which untreated control cultures reached the stationary phase, typically 12-15 h, to compare growth within equivalent phases.

Protease dose-response curves were generated by fitting AUC values to a four-parameter logistic model in (v4.4.3) ^74^. IC20, IC50, and IC70 values were estimated from the fitted curves and represent the protease concentrations required to reduce bacterial growth by 20%, 50%, or 70%, respectively, relative to untreated controls. For comparison of strain-specific protease sensitivity, a growth inhibition index (GI) was calculated from AUC values:

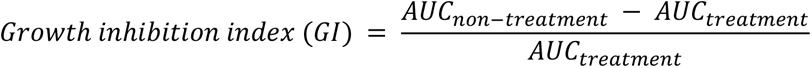

where AUCnon-treatment and AUCtreatment represent the AUC values of untreated and protease-treated cultures, respectively. Statistical differences in GI between strains were assessed using two-sided Mann-Whitney U tests in Python (v3.9). Data from three independent experiments were pooled within predefined low- and high-inhibition categories, corresponding approximately to IC20-IC40 and IC50-IC80, respectively. Low- and high-inhibition ranges were defined as 0.01-0.08 and 0.1-0.8 mg/mL for trypsin, and 0.001-0.008 and 0.01-0.06 mg/mL for proteinase K.

### Antimicrobial Peptide sensitivity assay

*P. vulgatus^ATCC^* ^8482^ wildtype and the isogenic *Δdpp11a* mutant were grown to mid-exponential phase in reinforced clostridial medium (RCM, 37.5 g L⁻¹, VWR) under anaerobic conditions (85% N₂, 10% H₂, 5% CO₂, 0% O₂). Cells were harvested by centrifugation (800×*g*, 10 min, 4°C), washed, and resuspended in ice-cold 10 mM sodium phosphate buffer (pH 7.4). Approximately 1×10^7^ colony-forming units (CFU) were incorporated into an underlay gel consisting of 0.1% EEO agarose (Sigma) and 0.1% RCM in 10 mM sodium phosphate buffer (pH 7.4). Upon solidification, 2.5-mm-diameter wells were punched into the gel and loaded with 1 µg of antimicrobial peptides (hBD-2, hBD-3, and LL-37; 0.25 µg/µL). Acetic acid (0.01%) and 1 µg of lysozyme (0.25 µg/µL) were utilized as negative and positive controls, respectively. The underlay gel was incubated for 3 h to allow the diffusion of the antimicrobial peptides into the gel. Subsequently, an overlay gel composed of 3% RCM, 0.1% EEO agarose, and 10 mM sodium phosphate buffer (pH 7.4) was applied. Plates were incubated at 37°C under anaerobic conditions for 24 h, and antimicrobial activity was quantified by measuring the zone of inhibition. All assays were performed in biological triplicates.

### Phage isolation

Sewage water collected from multiple locations in Munich were centrifuged (6,000 x *g*, 10 min) and sterile-filtered (0.22 μm). Sterile filtrates were enriched with *P. vulgatus* host strains by overnight anaerobic incubation in a 1:1 mixture of sewage filtrate and 2x BHI medium containing a 1:100 dilution of an overnight bacterial culture. Phage lysates were recovered by centrifugation and filtration and screened by spotting onto double-layer agar overlays seeded with the corresponding host strain. Individual plaques were isolated, propagated on the respective host strain, and subjected to three rounds of plaque purification before preparation of clonal phage lysates, which were stored at 4°C.

### Spot Assay

Overnight bacterial cultures were diluted 1:40 in fresh mGAM medium and grown anaerobically at 37°C to mid-log phase (OD600nm = 0.3-0.5). Aliquots (100 μL) were mixed with 4 mL molten top agar (0.7%) and overlaid onto pre-warmed mGAM agar plates. Phage stocks were serially diluted 10-fold in Saline Magnesium (SM) buffer (200 mM NaCl, 10 mM MgSO4, 50 mM Tris-HCl, pH 7.5), and 10 μL of each dilution was spotted onto the bacterial lawn. Plates were incubated anaerobically at 37°C for 24-48 h. Zones of clearing were scored as evidence of phage susceptibility, and phage titers were determined as plaque-forming units (PFU/mL).

### Phage Amplification

Phage stocks were amplified by infecting exponentially growing *P. vulgatus^H^*^253^ wildtype cultures (1:100 dilution of overnight culture in mGAM) with 10 μL phage lysate and incubating anaerobically at 37°C for 16 h. Phage lysates were recovered by centrifugation and filtration (0.2 μm), treated with 10% (v/v) chloroform to remove residual bacterial contaminants, and stored at 4°C. Phage titers were determined by spot assay as described above. When required, high-titer phage stocks were generated by concentrating lysates and exchanging the buffer to SM using 100-kDa MWCO centrifugal filter units (Amicon Ultra, Merck Millipore).

### Phage genome sequencing

#### Phage genomic DNA extraction

Phage genomic DNA was extracted from clarified phage lysates following DNase and RNase treatment to remove contaminating host nucleic acids. Viral capsids were lysed by proteinase K/SDS digestion, and genomic DNA was purified by phenol-chloroform extraction, ethanol precipitation, and column-based cleanup (Macherey-Nagel). DNA quality was verified by agarose gel electrophoresis before sequencing.

#### Genome sequencing, assembly, and annotation

Phage genomes were sequenced on an Illumina MiSeq Nano platform (2 × 150 bp). Reads were quality assessed using FastQC and trimmed with Trimmomatic before de novo assembly with SPAdes ^75,76^. Viral contigs were identified using ViralVerify, and genome completeness was assessed with CheckV; only assemblies with >90% estimated completeness were retained for downstream analyses ^77^. Gene prediction and functional annotation were performed using geNomad ^78^.

#### Phage infection growth assay

Phage susceptibility was assessed using a 96-well kinetic growth assay. Overnight cultures of *P. vulgatus^H^*^253^ wildtype and the isogenic *Δdpp11a* mutant were diluted in mGAM to an initial OD600nm of 0.1 and infected with AvS7 or Lim7 across a dilution series corresponding to MOIs of 1 to 10^-6^. Infection kinetics across this MOI range were used to select 10^-4^ as the standardized MOI for subsequent strain-comparison experiments.

For strain comparisons, *P. vulgatus^H^*^253^ wildtype, isogenic *Δdpp11a*, complemented *Δdpp11a*::pNBU2::*dpp11a*, and catalytically inactive *Δdpp11a*::pNBU2::*dpp11a^S^*^656^*^N^* strains were infected with AvS7 or Lim7 at an MOI of 10^-4^. Plate incubation, OD600nm monitoring, AUC calculation, and growth inhibition analysis were performed as described for the protease sensitivity assay. Data from three independent experiments were pooled, and statistical comparisons between strains were performed using two-sided Mann-Whitney U tests in Python (v3.9).

### HuMiX-Based Co-culture of Caco-2 Cells with *P. vulgatus* and phage treatment

#### Caco-2 cell culture

Caco-2 cells (DSMZ: ACC169, RRID: CVCL_0025) were maintained in high-glucose DMEM with GlutaMAX (Thermo Fisher Scientific, cat. no. 61965-026) supplemented with 10% heat-inactivated fetal bovine serum (FBS; Life Technologies, cat. no. A5256801). Cells were cultured at 37°C in a humidified 5% CO2 incubator. Medium was replaced every 2 days, and cells were passaged approximately once per week, depending on confluency.

#### HuMiX device preparation and Caco-2 cell seeding

Human-microbial co-culture experiments were performed using the Human-Microbial Crosstalk (HuMiX) device as previously described, with minor modifications ^79,80^. Devices contained separate nitrogen gas-flushed, microbial cell culture, epithelial cell culture, and perfusion chambers, delineated by silicone gaskets and separated by semipermeable membranes. The nitrogen chamber was continuously perfused with N2 at 0.1 L/h to maintain an oxygen gradient compatible with the co-culture of anaerobic bacteria and human epithelial cells. For the epithelial cell culture chamber, gaskets with ethyleneterephtalate (PET) membranes with 1 μm pores (it4ip, 2000M12/620m103/200250) were sterilized and coated with 50 μg/mL rat tail collagen (Sigma-Aldrich,cat. no. 122-20) for 3 h at 37°C. For the microbial cell culture chamber, gaskets with PET membranes with 50 nm pores (it4ip, 2000M12/860N051/200250) were coated with 0.025 mg/mL porcine gastric mucin (Sigma-Aldrich, cat. no. M1778-10G)for 1 h at 37°C. After removal of excess coating solution, membranes were air-dried under sterile conditions, and devices were assembled.

Caco-2 cells were seeded into the epithelial chamber by injecting 1.5 mL of cell suspension at 1.25 x 10^6^ cells/mL. Devices were incubated under static conditions for 2 h to allow cell attachment. Medium perfusion was then initiated through the perfusion and microbial chambers using DMEM supplemented with 10% FBS, while the epithelial chamber remained closed for the remainder of the experiment.

#### Bacterial inoculation and phage treatment in HuMiX

At day 6 after Caco-2 seeding, HuMiX devices were transitioned to serum-free culture conditions before bacterial inoculation to minimize potential interference of serum-derived proteins with bacterial protease activity. The epithelial and perfusion chambers were supplied with serum-free DMEM, whereas the microbial chamber was supplied with a 1:1 mixture of mGAM and DMEM.

*P. vulgatus^H^*^253^ wildtype and the isogenic *Δdpp11a* mutant were grown anaerobically in mGAM, adjusted to an OD600nmof 0.5 in 1:1 mGAM:DMEM, and inoculated into the microbial chamber. For phage-treated conditions, AvS7 was added at the time of bacterial inoculation at an MOI of 10⁻¹. After inoculation, devices were maintained under static conditions for 4 h to facilitate bacterial association with the mucin-coated membrane before perfusion was resumed.

#### Sample collection and processing

After 24 h of co-culture, HuMiX devices were disassembled, and mammalian and microbial gaskets were separated. Cell-covered membranes from the epithelial chamber were divided into two parts: one half was used for cell viability assessment, and the other half was processed for ZO-1 immunostaining.

For viability analysis, membranes were washed with PBS and incubated with TrypLE (Life Technologies, cat. no. 12605010) at 37°C and 5% CO2 for 7 min to detach epithelial cells. Detachment was stopped with DMEM containing 10% FBS, and cells were released by gentle pipetting. Cell suspensions were centrifuged at 300 x *g* for 3 min at 4°C, resuspended in PBS, and mixed with Trypan Blue (Merck, cat. no. T8154) for counting and assessment of Trypan Blue exclusion to indicate cell viability.

Microbial chamber contents were collected and centrifuged at 8,000 x *g* for 5 min at 4°C to separate bacterial pellets and supernatants. Pellets were resuspended in PBS for OD600nm measurement.

#### ZO-1 immunofluorescence staining

After HuMiX disassembly, cell-covered membranes were washed twice with PBS, fixed with 4% paraformaldehyde for 15 min at room temperature, washed again, and permeabilized with 0.1% Triton X-100 in PBS for 30 min. Membranes were blocked with 4% BSA in PBS for 1 h at room temperature and incubated with Alexa Fluor 488-conjugated anti-ZO-1 antibody (1:100 in 1% BSA; Thermo Fisher Scientific, cat. no. 33-9100; 1:100 in 1% BSA/PBS) under light-protected conditions. Membranes were then washed with PBS containing 0.1% Tween-20 and PBS, mounted cell-side down with DAPI-containing Fluoroshield mounting medium (Fluoroshield^TM^ mit DAPI, Sigma-Aldrich, cat. no. F6057-20mL) and stored at 4°C in the dark until imaging.

#### Confocal microscopy

ZO-1 and DAPI signals were imaged using a Zeiss LSM confocal microscope equipped with an Airyscan detector and a LD LCI Plan-Apochromat 25x/0.8 NA objective. Alexa Fluor 488 and DAPI were acquired sequentially using 488 nm and 405 nm excitation, respectively. Images were collected as 16-bit Z-stacks with a 0.45 μm step size and processed by 3D Airyscan reconstruction. Maximum-intensity projections were generated in FIJI for downstream analysis. Laser power, detector gain, scan settings, and Airyscan processing parameters were kept constant across samples.

#### Quantification of ZO-1 tight junction signal and nuclei shape index

ZO-1 signal was quantified from maximum-intensity projections using a custom MATLAB R2024b workflow. DAPI and ZO-1 channels were extracted from merged TIFF files and processed independently. Nuclei were segmented from the DAPI channel, and the resulting nuclear masks were used to estimate cell number, assess nuclear morphology, and define cell-associated regions. Nuclear morphology was quantified by measuring nuclear area, perimeter, eccentricity, solidity, and major and minor axis length. A nuclear shape index was calculated for each segmented nucleus as perimeter divided by the square root of nuclear area.

ZO-1 images were contrast-enhanced, smoothed, and thresholded to generate binary ZO-1 masks. To restrict analysis to cell-associated areas, nuclear masks were morphologically dilated, and ZO-1 signal outside this region was excluded. ZO-1-positive pixels were identified from the binary mask, whereas intensity values were extracted from the original ZO-1 image. For each image, total ZO-1 fluorescence intensity, cell-associated area, and ZO-1 density were quantified. ZO-1 density was calculated as the integrated ZO-1 fluorescence intensity within cell-associated ZO-1-positive pixels divided by the corresponding cell-associated area per field of view (FOV).

### Statistical analysis of ZO-1 density and nuclei shape index

ZO-1 fluorescence density and nuclear shape index were analyzed using Gaussian linear mixed-effects models to account for the hierarchical structure of the imaging data ^81^. Experimental condition was included as a fixed effect and biological replicate as a random intercept. Models were fitted by restricted maximum likelihood (REML).

ZO-1 fluorescence density was analyzed at FOV level using the model: ZO-1 density ∼ condition + (1 | biological replicate). Nuclear shape index was summarized as the median value of all segmented nuclei within each FOV and analyzed using the model: FOV median shape index ∼ condition + (1 | biological replicate). Fields of view were treated as repeated measurements nested within independent biological replicates

Statistical analysis was performed using a linear mixed-effects model with treatment as a fixed effect and biological replicate as a random effect as follows. Pairwise significance testing was performed using the Satterthwaite approximation for denominator degrees of freedom ^82,83^. Satterthwaite-adjusted p-values and 95% confidence intervals were calculated using the R packages lme4 ^84^ and pbkrtest ^83^. Model-estimated condition means are presented with 95% Wald confidence intervals derived from the fixed-effect variance-covariance matrix.

### Comparative analysis of protease distribution across bacterial species

Protease family composition was compared across 31 bacterial species selected from the available dataset (**Table S1**). Protease annotations were retrieved from the MEROPS database ^85^, and the number and relative abundance of each protease family were calculated for each species.

Annotated DPP sequences from the selected species were retrieved from MEROPS and subjected to all-against-all sequence similarity analysis using CLANS with default parameters to resolve sequence-based relationships among Dpp proteins ^86^. Sequence clusters were assigned using a stringent significance cutoff of p < 3.8 x 10^-25^, and more distant sequence relationships were displayed using an edge cutoff of 1 x 10^-6^. To compare the taxonomic distribution of Dpp subgroups, each species was scored for the presence or absence of each subgroup, and the resulting binary matrix was visualized by hierarchical clustering. AlphaFold-predicted structures of *P. vulgatus* DPP were aligned using FoldMason ^87^ implemented in the Foldseek web server using default parameters. The resulting guide tree was used to compare structure-based relationships among DPP family members.

### Meta-transcriptomic analysis of DPPs and phage transcript abundance

Meta-transcriptomic data and clinical metadata were obtained from the Inflammatory Bowel Disease Multi-omics Database ^42^ and grouped as non-IBD, Ulcerative Colitis (UC), and Crohn’s Disease (CD). Reads were aligned using bbsplit from BBMap v38.18 ^88^ with a minimum sequence identity threshold of 0.9. Reference targets included phage genomes (AvS1, AvS3, AvS7, Lim1, Lim3, and Lim7), *P. vulgatus^ATCC^* ^8482^ DPP genes annotated in MEROPS^33^. Raw read counts were normalized as reads per kilobase per million mapped reads (RPKM) to account for transcript length and sequencing depth.

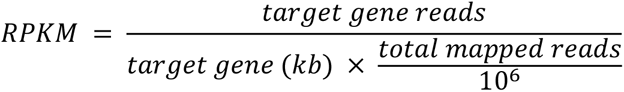

For each sample, phage transcript abundance was summarized as the mean RPKM across individual phage genomes with detectable reads per sample, generating a composite phage abundance measure for downstream analyses. Disease-associated differences in transcript abundance were assessed by comparing UC and CD samples separately against non-IBD controls using two-sided Mann-Whitney U tests in Python (v3.9).

## Supplementary Material

### Supplementary Content

#### Supplementary Figures

**Figure S1.**
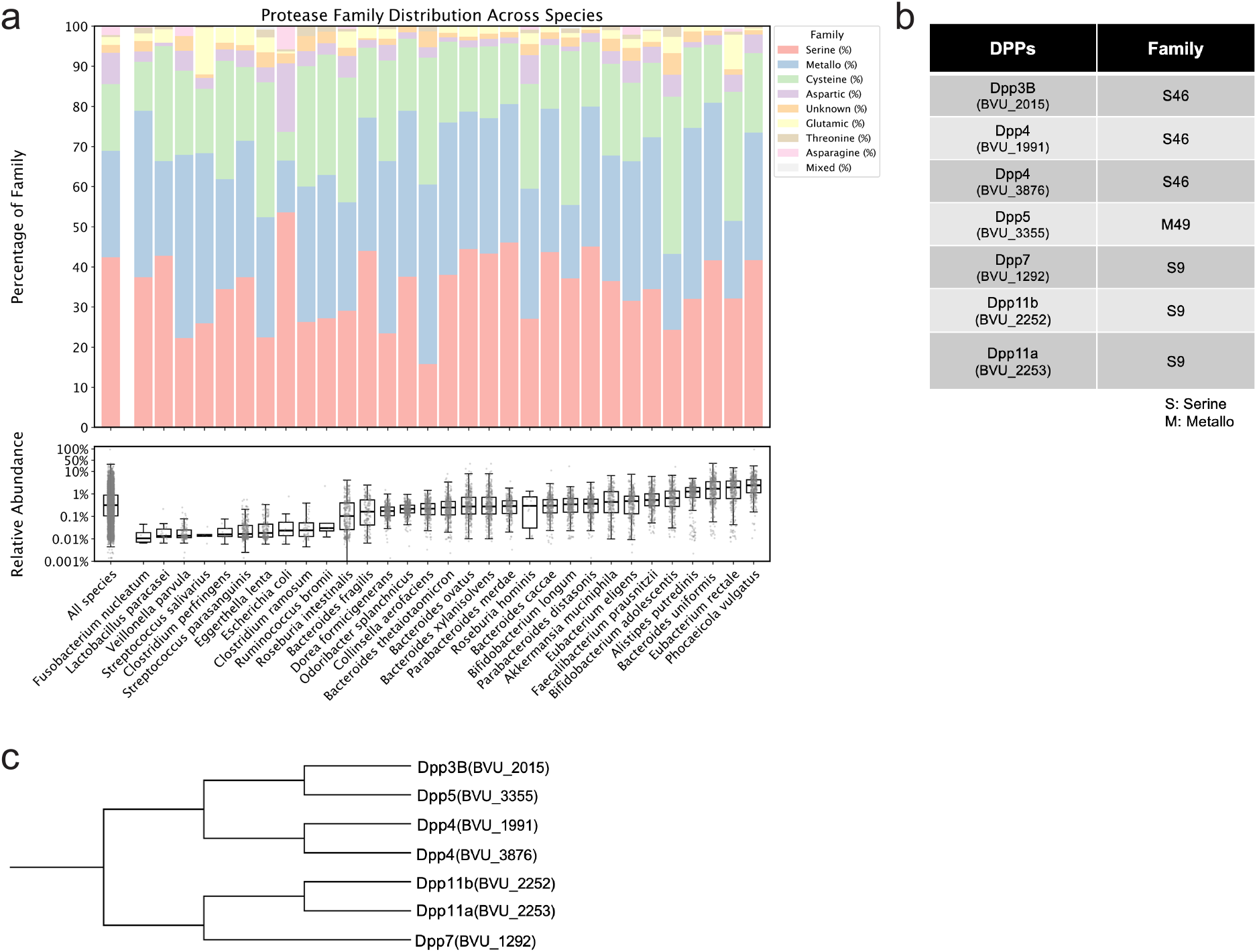
Protease encoding genes in gut bacteria, Dpp distribution and their structural similarity. **a**) Distribution of predicted protease families encoded by representative commensal gut bacterial genomes. Protease annotations were assigned using the MEROPS database^89^ and grouped according to catalytic family. Stacked bars (upper) show the proportion of annotated proteases belonging to each protease family for each species, expressed as a percentage of the total predicted proteases encoded by that genome. Species are ordered by increasing median relative abundance in healthy human gut metagenomes. The ‘All species column’ represents the combined protease family distribution across all analysed genomes. The lower panel shows the distribution of species relative abundance across healthy human metagenomic samples. Each point represents one individual sample, boxes indicate the interquartile range (IQR) with the center line denoting the median, and whiskers extend to the most extreme values within 1.5 × IQR. The ‘All species’ category summarizes the pooled relative abundances of all species included in the analysis. **b**) Table of *P. vulgatus*^ATCC 8482^ DPP’s and their protease family classification according to MEROPS^89^. **c**) Structural similarity of *P. vulgatus* DPP proteins. AlphaFold-predicted structures of DPP proteins were aligned using FoldMason in a structure-guided multiple structural alignment. The resulting guide tree depicts the relative structural similarity among Dpp proteins, with branch lengths corresponding to structural divergence inferred from the FoldMason alignment. Structures were predicted using AlphaFold and compared using FoldMason/Foldseek^87^.

**Figure S2.**
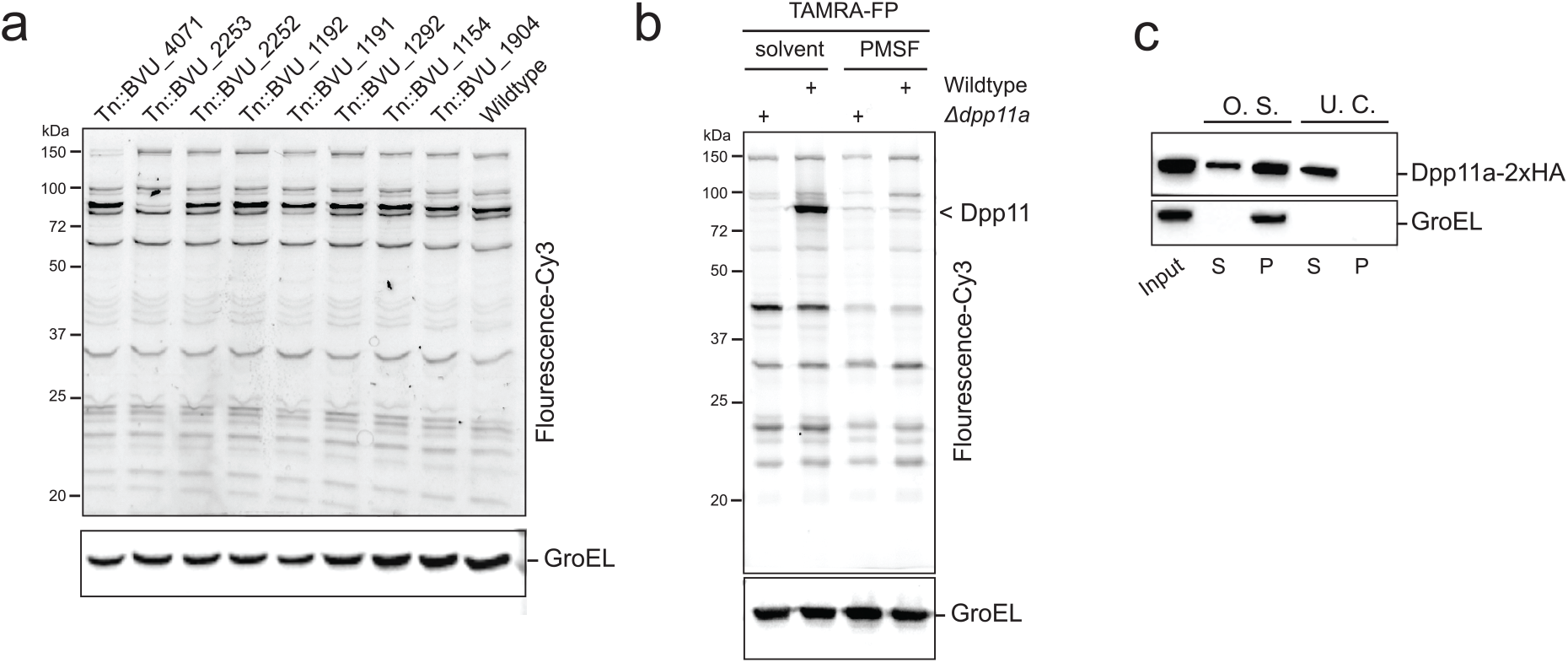
*P. vulgatus*^ATCC8482^ Dpp11 is an active serine hydrolase. **a**) Serine protease activity of cell pellets was assessed using TAMRA-FP on a panel of transposon insertion mutants. Experiment was performed and analysed as described in 2a. **b**) Serine protease activity was assessed on *P. vulgatus*^ATCC 8482^ in the presence of 1 µM Phenylmethylsulfonyl fluoride (PMSF) or solvent control. **c**) Osmotic Shock (O.S.) was performed on *P. vulgatus*^ATCC 8482^ expressing a chromosomal tandem c-terminally HA-tagged version of Dpp11 (Dpp11a-2xHA). Periplasmic proteins were separated from intact cells by low-speed centrifugation (S = supernatant, P = Pellet). Extracted periplasmic proteins were then subjected to ultracentrifugation (U. C.) to separate membrane-associated proteins extracted during O. S. from free-soluble extracted periplasmic proteins. Samples were analysed by immunoblot using anti-HA and anti-GroEL as a soluble protein control for cytoplasmic contamination. Representative data of three independent experiments (replicates in Supplementary Data).

**Figure S3.**
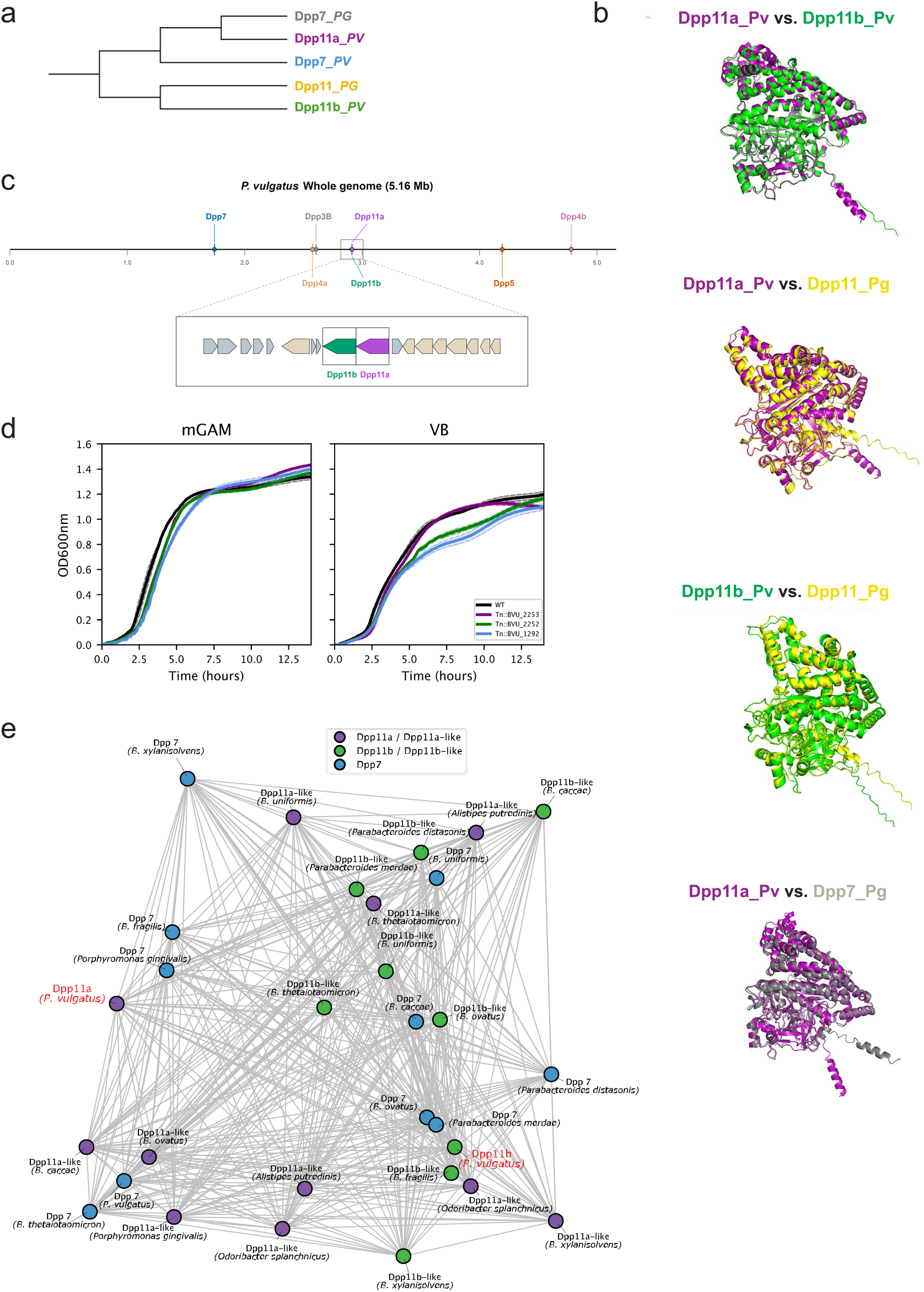
Structural relationships, genomic organization and functional divergence of Dpp11 paralogues in *P. vulgatus*. **a**) Structure-based clustering of Dpp proteins from *P. vulgatus* and *Porphyromonas gingivalis* (Pg). AlphaFold-predicted structures were compared by structure-guided multiple alignment using FoldMason^87^, and the resulting guide tree depicts the relative structural similarity among the proteins. Branch lengths represent distances derived from pairwise structural alignment scores. **b**) Structural superposition of *P. vulgatus* Dpp11 paralogues and *P. gingivalis* Dpp proteins. Shown are alignments of Dpp11a (purple) with Dpp11b (green), Dpp11a with *P. gingivalis* Dpp11 (yellow), Dpp11b with *P. gingivalis* Dpp11, and Dpp11a with *P. gingivalis* Dpp7 (grey). Structural alignments were performed using CEalign ^90^ in PyMOL. **c**) Genomic organization of DPP family genes within the *P. vulgatus* ATCC 8482 chromosome. The upper panel shows the chromosomal locations of DPP genes, and the lower panel highlights the *dpp11* locus, illustrating the adjacent arrangement of *dpp11a* and *dpp11b*. **d**) Growth of *P. vulgatus*^ATCC 8482^ Wild-type and transposon mutants disrupting *dpp11a*, *dpp11b*, or *dpp7* in nutrient-rich mGAM medium (left) or defined minimal VB medium (right). OD_600_ was monitored over time. Wild-type is shown in black, Tn::*dpp11a* in purple, Tn::*dpp11b* in green, and Tn::*dpp7* in blue. Lines represent the mean of technical replicates and shaded regions indicate ± s.e.m. Representative of three independent biological experiments. **e**) CLANS^86^ analysis of Dpp7-, Dpp11a-like-, and Dpp11b-like proteins identified among representative *Bacteroides* species. Nodes represent individual proteins and are coloured according to protein family (Dpp11a/Dpp11a-like, purple; Dpp11b/Dpp11b-like, green; Dpp7, blue). Grey edges connect proteins sharing sequence similarity above the clustering threshold. *P. vulgatus* Dpp11a and Dpp11b are highlighted in red.

**Figure S4.**
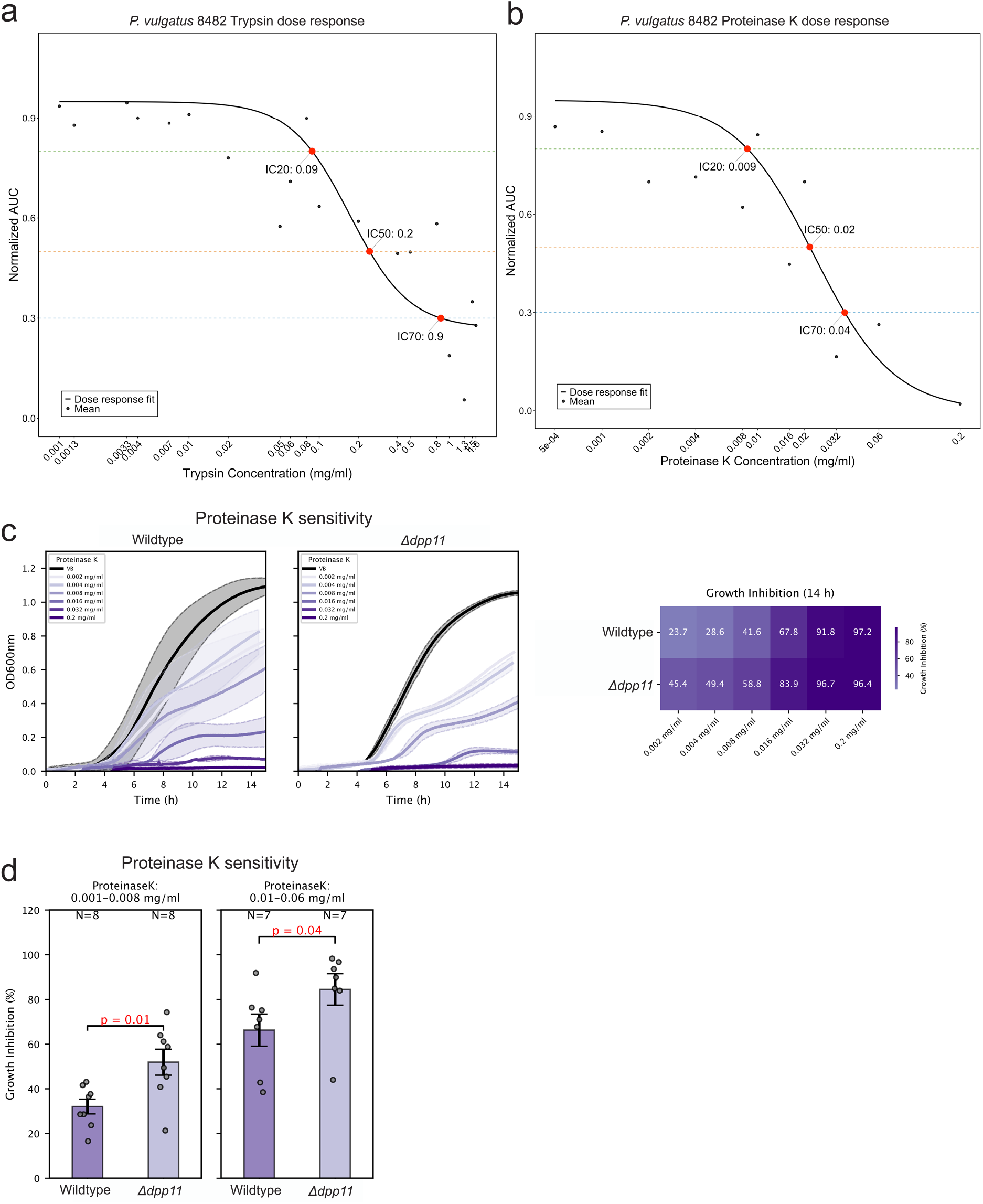
Dose-dependent sensitivity of *P. vulgatus* to extracellular serine proteases and the contribution of Dpp11 to protease resistance. **a)** dose-response analysis of *P. vulgatus*^ATCC 8482^ grown in semi-defined Varel-Bryant (VB) medium supplemented with increasing concentrations of porcine trypsin. Growth was quantified as the area under the growth curve (AUC) and normalized to untreated controls. Points represent mean normalized AUC values across seven independent experiments. The black line shows a four-parameter logistic dose-response curve fitted by nonlinear least-squares regression. Predicted IC20, IC50, and IC70 values, corresponding to 20%, 50%, and 70% growth inhibition, respectively, are indicated by red points. **b)** Dose-response analysis of *P. vulgatus* ATCC 8482 cultured in the presence of increasing concentrations of Proteinase K. Data are presented as in **a**. The fitted four-parameter logistic regression was used to estimate the IC20, IC50, and IC70 values. **c**) Representative growth curves of wild-type *P. vulgatus* ATCC 8482 and an isogenic Δ*dpp11* mutant cultured in increasing concentrations of Proteinase K (0.002-0.2 mg ml⁻¹) or untreated control (black). Technical replicates are shown as thin coloured lines and the mean as a thick line; shaded regions indicate ± s.d. The heatmap summarizes growth inhibition at 14 h, calculated from the area under the growth curve (AUC) relative to untreated controls. Representative of three independent experiments (replicates shown in Supplementary Data). d) Pooled Proteinase K sensitivity of wild-type and Δ*dpp11* strains at low (0.001-0.008 mg ml⁻¹) and high (0.01-0.06 mg ml⁻¹) protease concentrations. Growth inhibition (%) was calculated from the AUC relative to untreated controls. Bars represent mean ± s.e.m.; points represent individual measurements pooled from three independent experiments. *N* denotes the total number of measurements per genotype. Statistical significance was assessed using two-sided Mann-Whitney U test.

**Figure S5.**
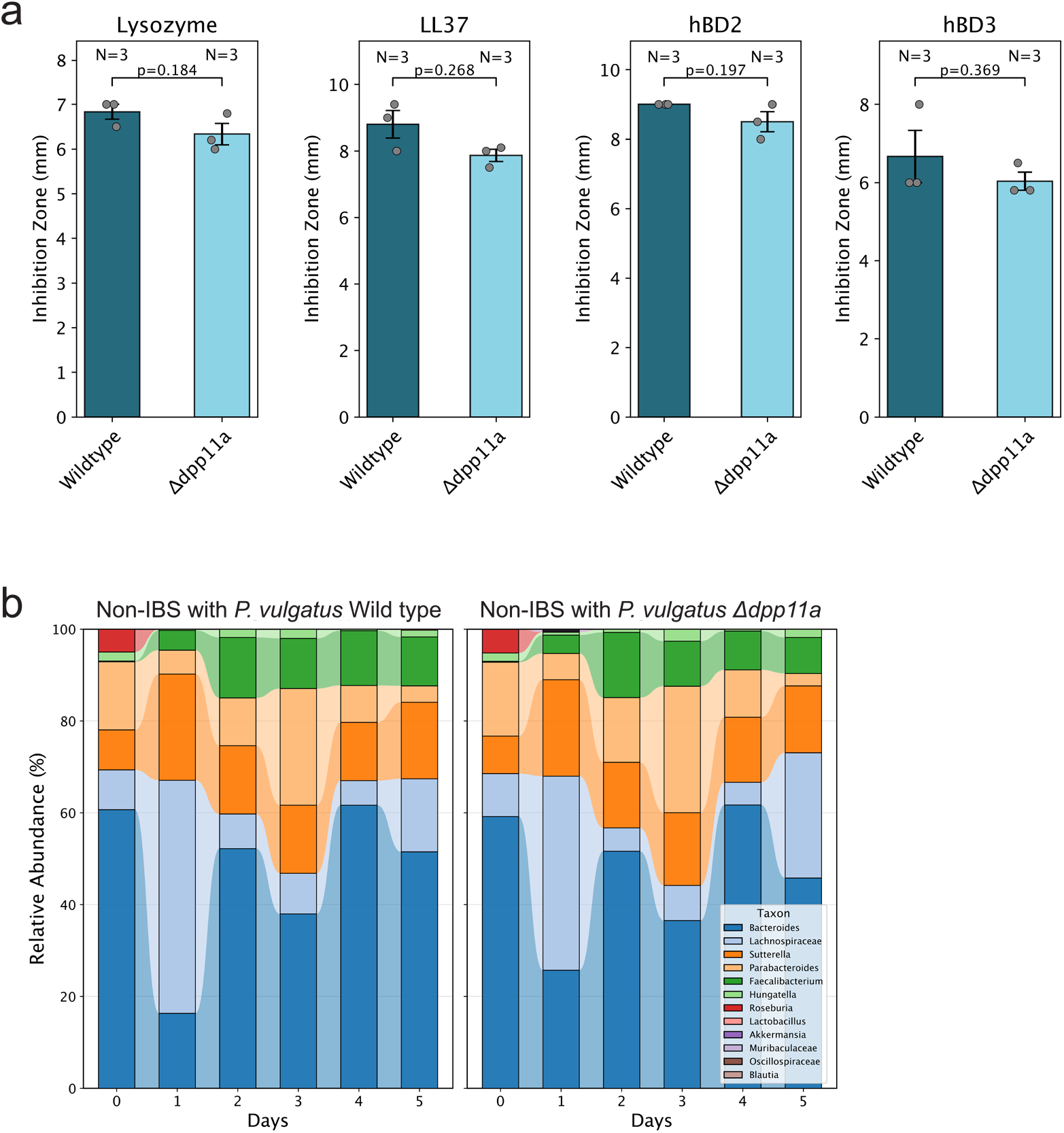
Dpp11a does not influence antimicrobial peptide susceptibility or synthetic gut community composition. **a**) Susceptibility of *P. vulgatus* ^ATCC 8482^ wild-type and Δ*dpp11a* strains to host antimicrobial proteins and peptides. Antimicrobial activity was assessed by agar diffusion assays using lysozyme, LL-37, human β-defensin 2 (hBD2), and human β-defensin 3 (hBD3). Inhibition zones (mm) are shown as mean ± s.e.m.; points represent independent biological replicates (N = 3). Statistical significance was assessed using a two-sided Mann-Whitney U test. Exact P values are indicated. **b**) Temporal composition of a defined non-IBD microbial community^68^ cultured in the presence of *P. vulgatus*^ATCC 8482^ wild-type (left) or the isogenic Δ*dpp11a* mutant (right). Community composition was determined by 16S rRNA gene amplicon sequencing over five days and is shown as the relative abundance of each bacterial genus. Data represent the average three independent biological experiments.

**Figure S6.**
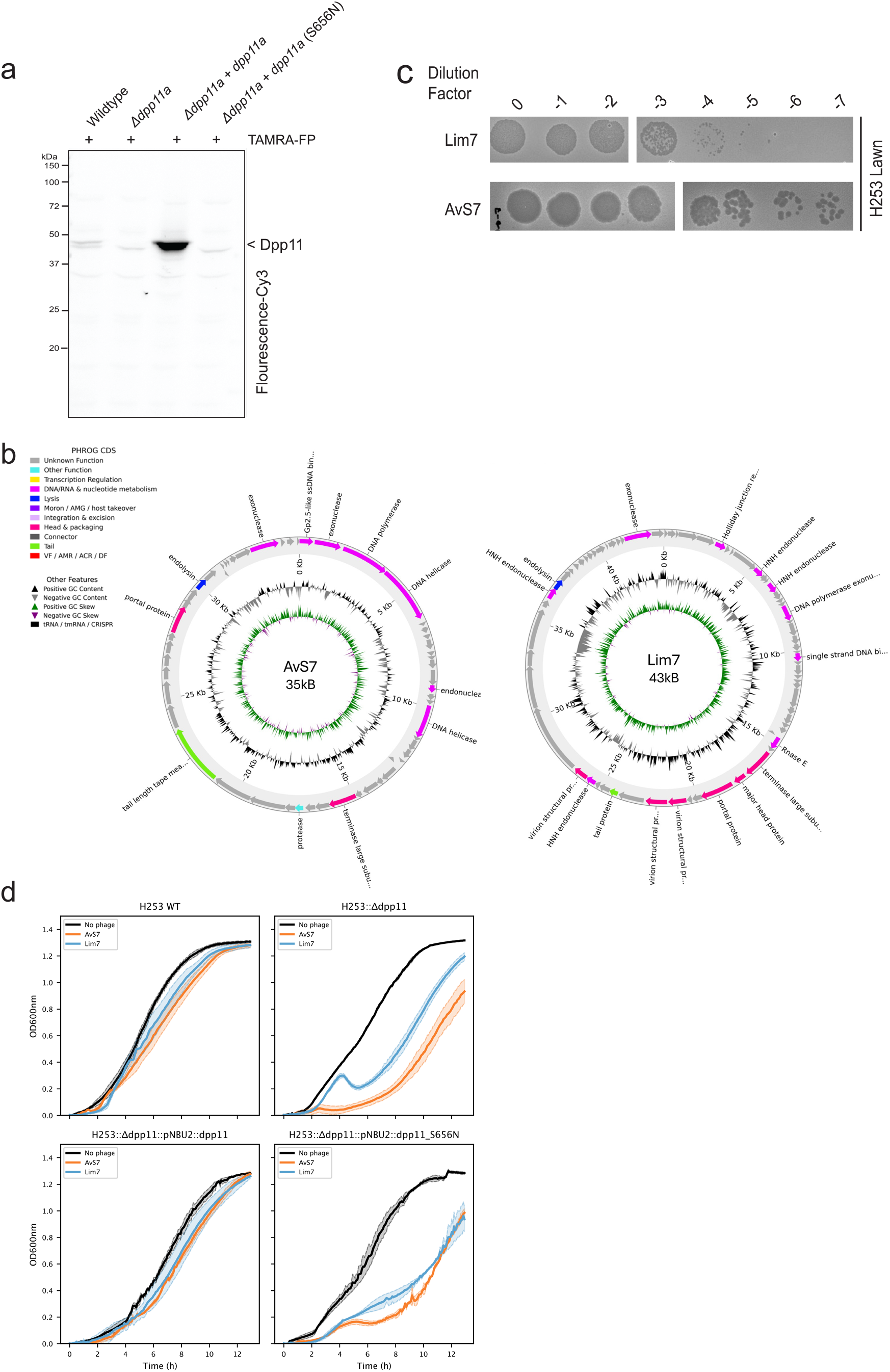
Verification of Dpp11a catalytic activity and characterization of *P. vulgatus*^H253^ bacteriophages. **a**) Activity-based protein profiling of Dpp11a complementation strains. Cell lysates from *P. vulgatus*^H253^ Wild type, the isogenic Δ*dpp11a* mutant, a chromosomally complemented strain expressing wild-type *dpp11a* (Δ*dpp11a*::pNBU2-*dpp11a*), and a complemented strain expressing the catalytic mutant Dpp11a(S656N) were labelled with the serine hydrolase activity-based probe TAMRA-FP and analysed by SDS-PAGE fluorescence scanning. Catalytic activity is restored by complementation with wild-type *dpp11a* but not by the S656N catalytic mutant. **b**) Circular genome maps of *P. vulgatus*^H253^ bacteriophages AvS7 (35 kb) and Lim7 (43 kb). Open reading frames (ORFs) are shown as arrows indicating gene orientation and are coloured according to predicted functional category. Inner rings depict GC content and GC skew. Functional annotations were assigned based on sequence homology and conserved domain analysis using the prokka package^91^. **c**) Plaque assays demonstrating infectivity of bacteriophages Lim7 and AvS7 on *P. vulgatus*^H253^. Ten-fold serial dilutions of phage lysates were spotted onto bacterial lawns, with dilution factors indicated above each spot. Plaque formation demonstrates productive infection and lysis of the bacterial host. **d**) Representative growth curves of *P. vulgatus*^H253^ Wild type, Δ*dpp11a*, the complemented strain (Δ*dpp11a*::pNBU2-*dpp11a*), and the catalytic mutant (Δ*dpp11a*::pNBU2-*dpp11a*(S656N)) following infection with AvS7 (orange) or Lim7 (blue). Uninfected controls are shown in black. Technical replicates are shown as thin coloured lines and the mean as a thick line; shaded regions indicate ± s.e.m. Representative of three independent biological experiments (replicates shown in Supplementary Data).

**Figure S7.**
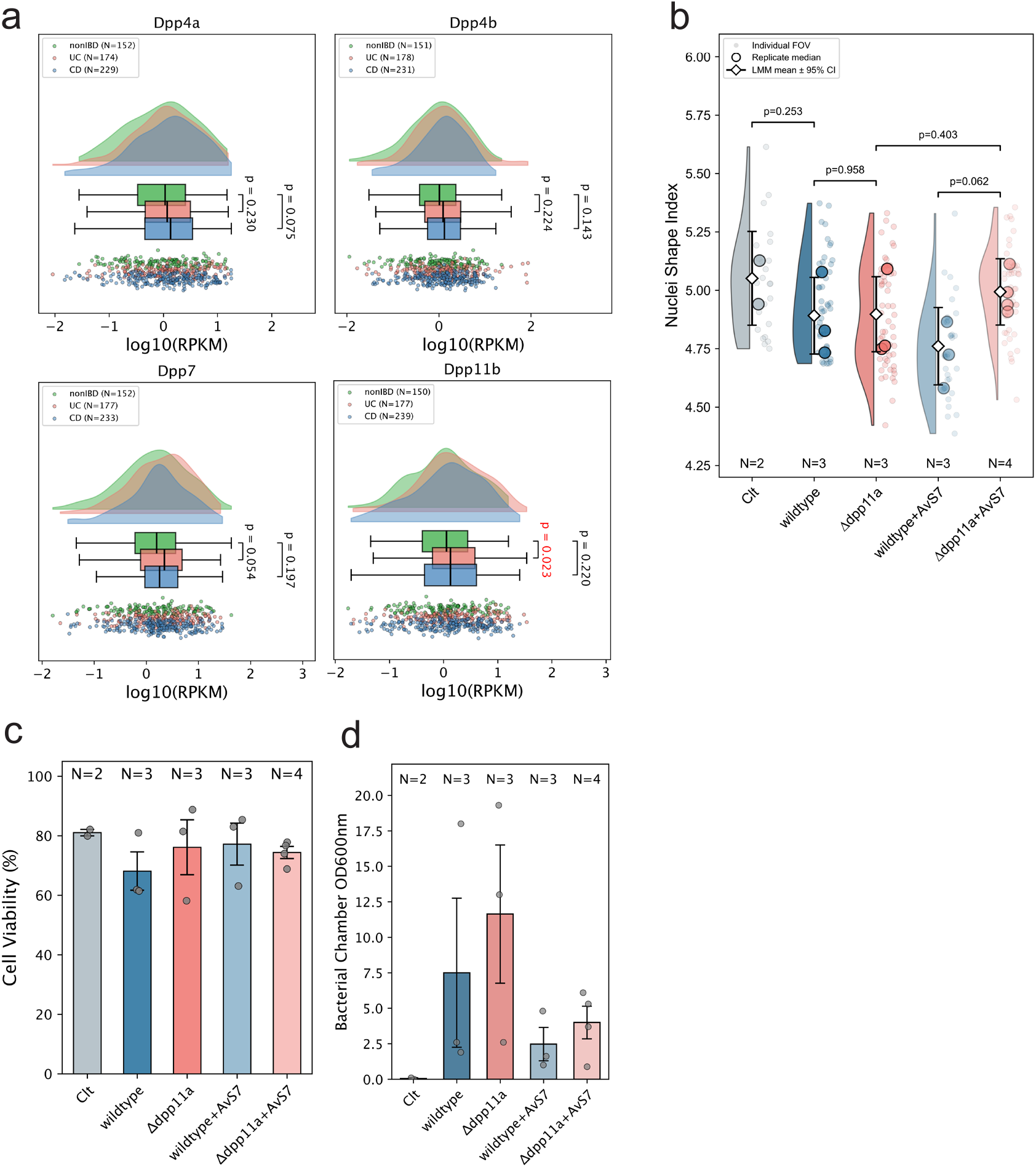
Dpp11 locus expression and epithelial responses are not accompanied by changes in nuclear morphology, epithelial viability, or bacterial abundance. **a)** Expression of *P. vulgatus dpp4a*, *dpp4b*, *dpp7*, and *dpp11b* in metatranscriptomic datasets from healthy non-IBD controls, patients with UC, and patients with Crohn’s disease (CD). Violin plots show the distribution of transcript abundance (log10 RPKM). Boxes indicate the interquartile range (IQR) with the center line denoting the median, whiskers extend to the most extreme values within 1.5 × IQR, and points represent individual samples. Statistical significance was assessed using two-sided Mann-Whitney U tests. **b)** Quantification of epithelial nuclear shape index from the HuMiX experiments corresponding to Fig. 5. Nuclear shape index was calculated for segmented nuclei and summarized as the median value for each field of view (FOV). Light points represent individual FOVs, large circles denote biological replicate medians, and diamonds indicate estimated marginal means ± 95% Wald confidence intervals from a Gaussian linear mixed-effects model. Experimental condition was included as a fixed effect and biological replicate as a random intercept. Pairwise comparisons were performed using the Satterthwaite approximation for denominator degrees of freedom. Exact *P* values are shown. *N* indicates the number of independent biological replicates. **c)** End point viability of Caco-2 epithelial cells following co-culture with the indicated *P. vulgatus* strains and phage treatments in the HuMiX system. Bars represent mean ± s.e.m., and points represent independent biological replicates. *N* indicates the number of biological replicates. **d)** End point bacterial abundance measurement within the microbial chamber of the HuMiX device, measured as optical density at 600 nm (OD_600_) at the conclusion of the experiment. Bars represent mean ± s.e.m., and points represent independent biological replicates. *N* indicates the number of biological replicates.

#### Supplementary Tables

**Table S1.** Strains used for protease distribution analysis

**Table S2.** Antibodies used in this study

**Table S3.** Software used in this study

**Table S4.** Strains used in this study

**Table S5.** Primers used in this study

**Table S6.** PCR conditions used in this study

## References

1. Antalis, T. M., Shea-Donohue, T., Vogel, S. N., Sears, C. & Fasano, A. Mechanisms of disease: protease functions in intestinal mucosal pathobiology. Nat. Clin. Pract. Gastroenterol. Hepatol. 4, 393–402 (2007).

2. Van Spaendonk, H. et al. Regulation of intestinal permeability: The role of proteases. World J. Gastroenterol. 23, 2106–2123 (2017).

3. Kiela, P. R. & Ghishan, F. K. Physiology of intestinal absorption and secretion. Best Pract. Res. Clin. Gastroenterol. 30, 145–159 (2016).

4. Caminero, A., Guzman, M., Libertucci, J. & Lomax, A. E. The emerging roles of bacterial proteases in intestinal diseases. Gut Microbes 15, 2181922 (2023).

5. Steck, N., Mueller, K., Schemann, M. & Haller, D. Bacterial proteases in IBD and IBS. Gut 61, 1610–1618 (2012).

6. Deraison, C. & Vergnolle, N. Proteases in intestinal health and disease. Nat. Rev. Gastroenterol. Hepatol. 23, 6–28 (2026).

7. Kriaa, A. et al. Serine proteases at the cutting edge of IBD: Focus on gastrointestinal inflammation. FASEB J. 34, 7270–7282 (2020).

8. Rolland-Fourcade, C. et al. Epithelial expression and function of trypsin-3 in irritable bowel syndrome. Gut 66, 1767–1778 (2017).

9 . Solà Tapias, N., et al. Colitis linked to endoplasmic reticulum stress induces trypsin activity affecting epithelial functions. J. Crohns. Colitis 15, 1528–1541 (2021).

10. Ducroc, R. et al. Trypsin is produced by and activates protease-activated receptor-2 in human cancer colon cells: evidence for new autocrine loop. Life Sci. 70, 1359–1367 (2002).

11. Santiago, A. et al. Crohn’s disease proteolytic microbiota enhances inflammation through PAR2 pathway in gnotobiotic mice. Gut Microbes 15, 2205425 (2023).

12. Porras, A. M. et al. Inflammatory Bowel Disease-Associated Gut Commensals Degrade Components of the Extracellular Matrix. MBio 13, e0220122 (2022).

13. Mills, R. H. et al. Multi-omics analyses of the ulcerative colitis gut microbiome link Bacteroides vulgatus proteases with disease severity. Nature Microbiology 1–15 (2022).

14. Lakemeyer, M. et al. A Bacteroides fragilis protease activates host PAR2 to induce intestinal pain and inflammation. Cell Host Microbe 33, 1686–1702.e11 (2025).

15. Jablaoui, A. et al. Fecal Serine Protease Profiling in Inflammatory Bowel Diseases. Front. Cell. Infect. Microbiol. 10, 21 (2020).

16. Norman, J. M. et al. Disease-specific alterations in the enteric virome in inflammatory bowel disease. Cell 160, 447–460 (2015).

17. Shkoporov, A. N. & Hill, C. Bacteriophages of the human gut: The ‘known unknown’ of the microbiome. Cell Host Microbe 25, 195–209 (2019).

18. Campbell, D. E. et al. Infection with Bacteroides Phage BV01 Alters the Host Transcriptome and Bile Acid Metabolism in a Common Human Gut Microbe. Cell Rep. 32, 108142 (2020).

19. Zuo, T. et al. Gut mucosal virome alterations in ulcerative colitis. Gut 68, 1169–1179 (2019).

20. Sinha, A. et al. Transplantation of bacteriophages from ulcerative colitis patients shifts the gut bacteriome and exacerbates the severity of DSS colitis. Microbiome 10, 105 (2022).

21. Majzoub, M. E. et al. The phageome of patients with ulcerative colitis treated with donor fecal microbiota reveals markers associated with disease remission. Nat. Commun. 15, 8979 (2024).

22. Clooney, A. G. et al. Whole-virome analysis sheds light on viral dark matter in inflammatory bowel disease. Cell Host Microbe 26, 764–778.e5 (2019).

23. Sato, K. et al. OmpA variants affecting the adherence of ulcerative colitis-derived Bacteroides vulgatus. J. Med. Dent. Sci. 57, 55–64 (2010).

24. Rath, H. C., Wilson, K. H. & Sartor, R. B. Differential induction of colitis and gastritis in HLA-B27 transgenic rats selectively colonized with Bacteroides vulgatus or Escherichia coli. Infect. Immun. 67, 2969–2974 (1999).

25. Hoentjen, F. et al. CD4(+) T lymphocytes mediate colitis in HLA-B27 transgenic rats monoassociated with nonpathogenic Bacteroides vulgatus. Inflamm. Bowel Dis. 13, 317–324 (2007).

26. Rath, H. C. et al. Normal luminal bacteria, especially Bacteroides species, mediate chronic colitis, gastritis, and arthritis in HLA-B27/human beta2 microglobulin transgenic rats. J. Clin. Invest. 98, 945–953 (1996).

27. Bloom, S. M. et al. Commensal Bacteroides species induce colitis in host-genotype-specific fashion in a mouse model of inflammatory bowel disease. Cell Host Microbe 9, 390–403 (2011).

28. Fernandes, M. A. et al. Enteric virome and bacterial Microbiota in children with ulcerative colitis and Crohn disease. J. Pediatr. Gastroenterol. Nutr. 68, 30–36 (2019).

29. Ohara-Nemoto, Y. et al. Asp- and Glu-specific novel dipeptidyl peptidase 11 of Porphyromonas gingivalis ensures utilization of proteinaceous energy sources. J. Biol. Chem. 286, 38115–38127 (2011).

30. Wang, K. et al. Microbial-host-isozyme analyses reveal microbial DPP4 as a potential antidiabetic target. Science 381, eadd5787 (2023).

31. Olivares, M. et al. Gut microbiota DPP4-like enzymes are increased in type-2 diabetes and contribute to incretin inactivation. Genome Biol. 25, 174 (2024).

32. Aljumaah, M. R., Roach, J., Hu, Y., Gunstad, J. & Azcarate-Peril, M. A. Microbial dipeptidyl peptidases of the S9B family as host-microbe isozymes. Sci. Adv. 11, eads5721 (2025).

33. Rawlings, N. D. & Bateman, A. How to use the MEROPS database and website to help understand peptidase specificity. Protein Sci. 30, 83–92 (2021).

34. Maier, L. et al. Extensive impact of non-antibiotic drugs on human gut bacteria. Nature 555, 623–628 (2018).

35. Zeevi, D. et al. Personalized nutrition by prediction of glycemic responses. Cell 163, 1079–1094 (2015).

36. Asnicar, F. et al. Microbiome connections with host metabolism and habitual diet from 1,098 deeply phenotyped individuals. Nat. Med. 27, 321–332 (2021).

37. Olsson, L. M. et al. Dynamics of the normal gut microbiota: A longitudinal one-year population study in Sweden. Cell Host Microbe 30, 726–739.e3 (2022).

38. Rigottier-Gois, L., Rochet, V., Garrec, N., Suau, A. & Doré, J. Enumeration of Bacteroides species in human faeces by fluorescent in situ hybridisation combined with flow cytometry using 16S rRNA probes. Syst. Appl. Microbiol. 26, 110–118 (2003).

39. Wang, S. et al. Metagenomic analysis of mother-infant gut microbiome reveals global distinct and shared microbial signatures. Gut Microbes 13, 1–24 (2021).

40. Jemielita, M., Mashruwala, A. A., Valastyan, J. S., Wingreen, N. S. & Bassler, B. L. Secreted proteases control the timing of aggregative community formation in Vibrio cholerae. MBio 12, e0151821 (2021).

41. Kriaa, A. et al. SP-1, a Serine Protease from the Gut Microbiota, Influences Colitis and Drives Intestinal Dysbiosis in Mice. Cells 10, (2021).

42. Lloyd-Price, J. et al. Multi-omics of the gut microbial ecosystem in inflammatory bowel diseases. Nature 569, 655–662 (2019).

43. Teufel, F. et al. SignalP 6.0 predicts all five types of signal peptides using protein language models. Nat. Biotechnol. 40, 1023–1025 (2022).

44. Nemoto, T. K. & Ohara Nemoto, Y. Dipeptidyl-peptidases: Key enzymes producing entry forms of extracellular proteins in asaccharolytic periodontopathic bacterium Porphyromonas gingivalis. Mol. Oral Microbiol. 36, 145–156 (2021).

45. Schimek, C. et al. Extraction of recombinant periplasmic proteins under industrially relevant process conditions: Selectivity and yield strongly depend on protein titer and methodology. Biotechnol. Prog. 36, e2999 (2020).

46. Li, Y. et al. Identification of trypsin-degrading commensals in the large intestine. Nature 609, 582–589 (2022).

47. Róka, R. et al. Colonic luminal proteases activate colonocyte proteinase-activated receptor-2 and regulate paracellular permeability in mice. Neurogastroenterol. Motil. 19, 57–65 (2007).

48. Ramare, F., Hautefort, I., Verhe, F., Raibaud, P. & Iovanna, J. Inactivation of tryptic activity by a human-derived strain of Bacteroides distasonis in the large intestines of gnotobiotic rats and mice. Appl. Environ. Microbiol. 62, 1434–1436 (1996).

49. Herbst, E. et al. Extracellular activity of a bacterial protease associated with reduced phage infectivity. PLoS One 21, e0332566 (2026).

50. Castillo, D. et al. Phage defense mechanisms and their genomic and phenotypic implications in the fish pathogen Vibrio anguillarum. FEMS Microbiol. Ecol. 95, (2019).

51. Górski, A., Borysowski, J. & Miȩdzybrodzki, R. Bacteriophage interactions with epithelial cells: Therapeutic implications. Front. Microbiol. 11, 631161 (2020).

52. Sutton, T. D. S. & Hill, C. Gut bacteriophage: Current understanding and challenges. Front. Endocrinol. (Lausanne) 10, 784 (2019).

53. Barr, J. J. Missing a phage: Unraveling tripartite symbioses within the human gut. mSystems 4, e00105–19 (2019).

54. Barr, J. J. et al. Bacteriophage adhering to mucus provide a non-host-derived immunity. Proc. Natl. Acad. Sci. U. S. A. 110, 10771–10776 (2013).

55. Kan, L. & Barr, J. J. A mammalian cell’s guide on how to process a bacteriophage. Annu. Rev. Virol. 10, 183–198 (2023).

56. Bichet, M. et al. Mammalian cells internalize bacteriophages and utilize them as a food source to enhance cellular growth and survival. bioRxiv 2023.03.10.532157 (2023) doi:10.1101/2023.03.10.532157.

57. Shah, P. et al. A microfluidics-based in vitro model of the gastrointestinal human-microbe interface. Nat. Commun. 7, 11535 (2016).

58. Hampton, H. G., Watson, B. N. J. & Fineran, P. C. The arms race between bacteria and their phage foes. Nature 577, 327–336 (2020).

59. Millman, A. et al. An expanded arsenal of immune systems that protect bacteria from phages. Cell Host Microbe 30, 1556–1569.e5 (2022).

60. Varel, V. H. & Bryant, M. P. Nutritional Features of Bacteroides fragilis subsp. fragilis. Appl. Microbiol. 28, 251–257 (1974).

61. Bobonis, J., et al. Bacterial retrons encode phage-defending tripartite toxin–antitoxin systems. Nature 609, 144–150 (2022).

62. García-Bayona, L. & Comstock, L. E. Streamlined Genetic Manipulation of Diverse Bacteroides and Parabacteroides Isolates from the Human Gut Microbiota. MBio 10, (2019).

63. Laemmli, U. K. Cleavage of structural proteins during the assembly of the head of bacteriophage T4. Nature 227, 680–685 (1970).

64. Robinson, M. D., McCarthy, D. J. & Smyth, G. K. edgeR: a Bioconductor package for differential expression analysis of digital gene expression data. Bioinformatics 26, 139–140 (2010).

65. Nossal, N. G. & Heppel, L. A. The release of enzymes by osmotic shock from Escherichia coli in exponential phase. J. Biol. Chem. 241, 3055–3062 (1966).

66. Jalalirad, R. Selective and efficient extraction of recombinant proteins from the periplasm of Escherichia coli using low concentrations of chemicals. J. Ind. Microbiol. Biotechnol. 40, 1117–1129 (2013).

67. Rathore, A. S., Bilbrey, R. E. & Steinmeyer, D. E. Optimization of an osmotic shock procedure for isolation of a protein product expressed in E. coli. Biotechnol. Prog. 19, 1541–1546 (2003).

68. Hitch, T. C. A. et al. Function-based selection of synthetic communities enables mechanistic microbiome studies. ISME J. 19, wraf209 (2025).

69. Godon, J. J., Zumstein, E., Dabert, P., Habouzit, F. & Moletta, R. Molecular microbial diversity of an anaerobic digestor as determined by small-subunit rDNA sequence analysis. Appl. Environ. Microbiol. 63, 2802–2813 (1997).

70. Berry, D., Ben Mahfoudh, K., Wagner, M. & Loy, A. Barcoded primers used in multiplex amplicon pyrosequencing bias amplification. Appl. Environ. Microbiol. 77, 7846–7849 (2011).

71. Klindworth, A. et al. Evaluation of general 16S ribosomal RNA gene PCR primers for classical and next-generation sequencing-based diversity studies. Nucleic Acids Res. 41, e1 (2013).

72. Lagkouvardos, I., Fischer, S., Kumar, N. & Clavel, T. Rhea: a transparent and modular R pipeline for microbial profiling based on 16S rRNA gene amplicons. PeerJ 5, e2836 (2017).

73. Edgar, R. C. UPARSE: highly accurate OTU sequences from microbial amplicon reads. Nat. Methods 10, 996–998 (2013).

74. Ritz, C., Baty, F., Streibig, J. C. & Gerhard, D. Dose-response analysis using R. PLoS One 10, e0146021 (2015).

75. Bankevich, A. et al. SPAdes: a new genome assembly algorithm and its applications to single-cell sequencing. J. Comput. Biol. 19, 455–477 (2012).

76. Bolger, A. M., Lohse, M. & Usadel, B. Trimmomatic: a flexible trimmer for Illumina sequence data. Bioinformatics 30, 2114–2120 (2014).

77. Nayfach, S., Shi, Z. J., Seshadri, R., Pollard, K. S. & Kyrpides, N. C. New insights from uncultivated genomes of the global human gut microbiome. Nature 568, 505–510 (2019).

78. Camargo, A. P. et al. Identification of mobile genetic elements with geNomad. Nat. Biotechnol. 42, 1303–1312 (2024).

79. Lucchetti, M. et al. An organ-on-chip platform for simulating drug metabolism along the gut-liver axis. Adv. Healthc. Mater. 13, e2303943 (2024).

80. De Rudder, C., et al. immunoHuMiX: A personalizable gut-on-chip model for unraveling human microbiome–immune interactions. View (Beijing) 20250213 (2026).

81. Gelman, A. & Hill, J. Data Analysis Using Regression and Multilevel/hierarchical Models. (Cambridge University Press, 2007).

82. Satterthwaite, F. E. An approximate distribution of estimates of variance components. Biometrics 2, 110–114 (1946).

83. Halekoh, U. & Højsgaard, S. A Kenward-Roger approximation and parametric bootstrap methods for tests in linear mixed models - TheRPackagepbkrtest. J. Stat. Softw. 59, 1–32 (2014).

84. Bates, D., Mächler, M., Bolker, B. & Walker, S. Fitting linear mixed-effects models Usinglme4. J. Stat. Softw. 67, 1–48 (2015).

85. Rawlings, N. D. et al. The MEROPS database of proteolytic enzymes, their substrates and inhibitors in 2017 and a comparison with peptidases in the PANTHER database. Nucleic Acids Res. 46, D624–D632 (2018).

86. Frickey, T. & Lupas, A. CLANS: a Java application for visualizing protein families based on pairwise similarity. Bioinformatics 20, 3702–3704 (2004).

87. Gilchrist, C. L. M., Mirdita, M. & Steinegger, M. Multiple protein structure alignment at scale with FoldMason. Science 391, 485–488 (2026).

88. Bushnell, B. BBMap: A Fast, Accurate, Splice-Aware Aligner. https://escholarship.org/uc/item/1h3515gn (2014).

89. Rawlings, N. D., O’Brien, E. & Barrett, A. J. MEROPS: the protease database. Nucleic Acids Res. 30, 343–346 (2002).

90. Shindyalov, I. N. & Bourne, P. E. Protein structure alignment by incremental combinatorial extension (CE) of the optimal path. Protein Eng. 11, 739–747 (1998).

91. Seemann, T. Prokka: rapid prokaryotic genome annotation. Bioinformatics 30

